# Diet-induced obesity results in endothelial cell desensitization to VEGF-A and permanent islet vascular dysfunction

**DOI:** 10.1101/2023.11.06.565915

**Authors:** Yan Xiong, Andrea Dicker, Montse Visa, Erwin Ilegems, Per-Olof Berggren

## Abstract

**Background:** Pancreatic islet microvasculature is essential for optimal islet function and glucose homeostasis. However, islet vessel pathogenesis and its role in the manifestation of metabolic disorders remain understudied. Here we depict a time-resolved decline of intra-islet endothelial cell sensitivity to vascular endothelial cell growth factor A (VEGF-A) in a mouse model of diet-induced obesity.

**Methods:** Mice were transplanted with reporter islets in their eyes and put on different diet schemes for 48 weeks. Islet vascular morphology, VEGF-A signaling activity in islet endothelial cells and vessel function were longitudinally monitored by in vivo imaging, while the metabolic implication of islet vessel alterations was measured by glucose tolerance tests and insulin secretion assays.

**Results:** In parallel with substantial islet vasculature remodeling, diminished VEGF-A response in islet endothelial cells emerged after 12 weeks of western diet feeding. This led to vessel barrier dysfunction and hemodynamic dysregulation, which delayed transportation of secreted insulin into the blood. Islet vessels also exhibited a remarkable metabolic memory long after the removal of western diet. Neither islet endothelial cell VEGF-A sensitivity nor the vascular damage elicited by 24 weeks of western diet feeding was restored by switching to control diet for another 24 weeks. As a result, these refed mice still exhibited mild but significant impairment in glucose clearance, despite a complete normalization of body weight and insulin sensitivity. While plasma levels of soluble VEGF receptor 1 – the natural VEGF-A trap – were similar in all diet groups, increased activity of atypical protein kinase C (aPKC) was observed under both western diet and recovery conditions, which inhibited VEGF receptor 2 (VEGFR2) internalization and dampened VEGF-A triggered signal transduction in vivo and in human endothelial cells cultured under diet-mimicking conditions.

**Conclusions:** Long-term western diet feeding causes irreversible VEGF-A desensitization in islet endothelial cells and islet vessel dysfunction which undermines glucose homeostasis.

## Introduction

As an integral part of pancreatic islets, the rich microvascular network not only ensures nutritional provision for islet survival, but also provides critical developmental cues during pancreatic organogenesis and modulates its endocrine function at later stages[1–3]. Islet endothelial cells lining the luminal surface of the vessels are highly fenestrated and serve as gateways for efficient substance exchange between the interstitial fluid of islets and the blood stream, which is imperative for the islet sensing of plasma nutrient fluctuations as well as for timely hormone outflow[4, 5]. Obesity and diabetes have proved to elicit morphological remodeling of islet architecture both in humans and rodent models, and suboptimal vessel density is detrimental to islet function and glucose metabolism[6–8]. While increased oxidative stress concomitant with these metabolic disorders also leads to metabolic reprogramming and functional modifications in endothelial cells[9, 10], due to technical limitations it remains so far unclear whether intra-islet vessel functionality is undermined in obesity, which would restrict islet response to glucose.

VEGF-A plays a fundamental and indispensable role in vessel physiology and homeostasis. Upon binding to its principal signaling receptor VEGFR2, it initiates rapid internalization of the activated receptor and subsequent phosphorylation of key downstream signaling molecules, thereby promoting angiogenesis, increasing vascular permeability, as well as participating in the tuning of vasomotor activity[11–13]. In pancreatic islets, VEGF-A is constitutively expressed by all types of endocrine cells and signals through the endothelial membrane–localized VEGFR2. This intra-islet paracrine signal regulates islet vascular growth and patterning as well as coordinates innervation during pancreas development[14–16]. Tightly controlled VEGF-A expression is not only essential for the maintenance of the dense and highly permeable islet microvasculature, but also crucial for islet adaptation to changes in metabolic demands in terms of insulin secretion and β cell mass[17–21]. Moreover, VEGF-A is required for re-establishing blood flow in transplanted islets, which is imperative for graft survival[14, 22]. We have previously shown in a ciliopathy mouse model that primary cilia defects lead to VEGF-A/VEGFR2 signaling disruption in endothelial cells and consequently delay re-vascularization of transplanted islets[23].

Recently, VEGF-A has also been recognised as an underlying factor for metabolic diseases[24, 25]. Elevated plasma VEGF-A concentrations were repeatedly reported in obese and diabetic individuals[26–28]. Disturbance of local VEGF-A expression level and its signaling activity has also been implicated in aberrant angiogenesis associated with diabetes in a tissue-specific manner[29–32]. Upregulated VEGF-A signaling in diabetic retinal vessels results in excessive angiogenesis and leads to vascular lesions[33], whereas insufficient VEGF-A signaling underlies impaired wound healing[34, 35]. In addition, a ligand-independent non-canonical VEGFR2 signaling pathway was reported in cultured endothelial cells that were exposed to high glucose concentrations, which reduced the availability of surface VEGFR2 and diminished VEGF-A–triggered signal transduction[36]. Here, by integrating state-of-the-art in vivo imaging techniques with multiple vessel function assays, we provide new evidence that pancreatic islet endothelial cells develop resistance against VEGF-A under long-term western diet feeding. Moreover, we demonstrated that dysregulated VEGFR2 internalization and obstructed downstream signaling underlies irreversible islet vessel functional impairments, which compromise the efficiency of islet insulin output and glucose homeostasis.

## Methods

### Animal models and diet intervention

C57BL/6J, *Ai38* (#014538) and *Cdh5-Cre* mouse strains (#017968) were obtained from the Jackson laboratory (Maine, USA), the latter two strains were both backcrossed into C57BL/6J genetic background. At 6-8 weeks of age, male animals were given control diet (#EF CD88137, Ssniff Spezialdiäten, Germany) when they received islet transplantation into their eyes. Animals were then divided randomly into CD and WD groups, the control group continued to receive control diet, while the WD group was given high sugar/high fat WD instead (#EF TD88137 mod, 0.21% cholesterol, Ssniff Spezialdiäten, Germany). Experimental procedures involving live animals were carried out in accordance with the Karolinska Institutet’s guidelines for the care and use of animals in research and were approved by the institute’s Animal Ethics Committee (Ethical permit numbers 19462-2017 and 17431-2021).

### Islet isolation and transplantation

Islets were isolated by cannulation of the common bile duct of donor animals and infusion of 3 ml collagenase P, 0.8 mg/ml in HBSS containing 25 mM HEPES and 0.2% BSA (Sigma-Aldrich, USA). Inflated pancreata were carefully dissected out and digested in 37°C water bath for 10 min, and individual islets were hand-picked and further cultured in RPMI-1640 Medium (11mM D-glucose) supplemented with 10% fetal bovine serum, 100 IU/ml penicillin, 100 μg/ml streptomycin and 2 mM L-Glutamine (all from Thermo Fisher Scientific, USA).

For transplantation, recipient mice were anesthetized with 2% isoflurane (Baxter, USA) and fixed with a custom-made head and eye holder (Narishige, Japan) as previously described[37]. A small incision was made in the cornea with a 25 G needle and the tip of a glass cannula containing islets was inserted through the opening into the anterior chamber of the eye. 4-5 islets (150-200 μm in diameter) were carefully positioned on the iris around the pupil. Post-operative analgesia was done by subcutaneous injection of 0.1 ml/kg Temgesic (RB Pharmaceuticals Limited, UK). For [Ca^2+^] imaging experiments, 3-4-month-old male C57BL/6J mice were used as islet donors, and islets were kept in culture for a week before transplantation. 6-8-week-old male EC-GC3 mice were used as recipients. For the other *in vivo* imaging experiments, 3-4-month-old and 6-8-week-old male C57BL/6J mice were used as islet donors and recipients respectively, and islets were transplanted after overnight culture.

### Glucose and insulin tolerance test

Animals were fasted for 6 hours in all experiments. For intraperitoneal glucose tolerance tests, an amount of 2 g/kg body weight D-glucose (20% solution in PBS+/+, Merck, USA) was injected. For intraperitoneal insulin tolerance tests, 0.25 U/kg insulin (diluted in saline, Novo Nordisk, Denmark) was injected. Blood was taken from tail veins and blood glucose levels were measured at specified time points using an Accu-Chek Aviva system (Roche, Switzerland). For intravenous glucose tolerance tests, animals were anesthetized with a mixture of 0.5 mg/kg fentanyl, 16 mg/kg fluanisone (Hypnorm, Roche, Switzerland) and 8 mg/kg midazolam (Hameln, Sweden) after fasting. They were then placed on a heating pad (37 °C) under a heating lamp, and oxygen was supplied during the experiments through nose masks at 250 ml/min. 0.5 g/kg D-glucose (10% solution in PBS+/+) was injected through the tail vein, and blood was collected for the measurement of both glucose (Accu-Check) and insulin levels (Insulin AlphaLISA detection kit, Perkin Elmer, USA).

### In vivo imaging of islet grafts

Imaging was performed at specified time points post-transplantation by confocal microscopy. Animals were anesthetized with either 2% isoflurane inhalation (Baxter, USA), or a mixture of 0.5 mg/kg fentanyl, 16 mg/kg fluanisone (Hypnorm, Roche, Switzerland) and 8 mg/kg midazolam (Hameln, Sweden). During imaging sessions, animals were kept on a heating pad (37 °C) with a head holder in the heated microscope chamber, and oxygen was supplied throughout the experiments at 250 ml/min. Images of islet grafts were obtained with Leica DM6000 CFS/TCS SP5 confocal microscope equipped with a 25× objective (N.A. 0.95, Leica Microsystems, Germany). Pre-warmed viscotears (Laboratoires Théa, France) was used as an immersion medium between the lens and the mouse eyes. Morphology of islet graft was visualized by obtaining backscatter signal in all the imaging sessions (Ex.: 633 nm, Em.:630-636 nm).

For the visualization of islet vasculature, 100 μl of 2.5 mg/ml of 40-kDa FITC-conjugated dextran (Thermo Fisher Scientific, USA) in PBS+/+ was injected intravenously prior to imaging. Z-stacks of 2 μm thickness per step were acquired for every islet graft (Ex.: 488 & 633 nm, Em.: 500-540 nm & 630-636 nm).

For [Ca^2+^] imaging, a tail vein catheter containing 100 IE/ml heparin and 10 μg/ml mouse VEGF-A in sterile saline was inserted after anesthesia and prior to imaging. 10 μl heparin was directly injected, while 100 μl/30 g VEGF-A was injected after 1 min of baseline signal collection. Z stacks of 140 μm thickness in total (4 μm per step) were acquired with resonance scanner for every islet graft (Ex.: 488 & 633 nm, Em.: 500-550 nm & 630-636 nm).

For quantification of vascular leakage, a tail vein catheter containing 100 IE/ml heparin and 2.5 mg/ml of 40 kDa FITC-conjugated dextran in PBS+/+ was inserted after anesthesia and prior to imaging. 10 μl heparin was directly injected, while 100 μl/ 30 g of FITC-dextran was injected 1 min later. Z stacks of 40 μm thickness in total (2 μm per step) were acquired with resonance scanner for every islet graft (Ex.: 488 & 633nm, Em.: 500-540 nm & 630-636 nm).

For monitoring blood flow dynamics, RBCs were labelled as previously described[38]. Briefly, 35 μl blood was extracted from the tail vein, and 70 μl binding buffer (128 mM NaCl, 15 mM glucose, 10 mM HEPES, 4.2 mM NaHCO_3_, 3 mM KCl, 2 mM MgCl_2_, and 1 mM KH_2_PO_4_, pH adjusted to 7.4) was added. RBCs were purified by centrifugation at 1500 rpm (220 x g) for 5 min and resuspended in 140 μl binding buffer, and further labelled with 3.5 μl carbocyanide 1,1′-dioctadecyl-3,3,3′,3′-tetramethylindodicarbocyanine, 4-chlorobenzenesulfonate salt (DiD solid, Thermo Fisher Scientific, USA) for 10 min at 37 °C. The fluorescent RBCs were then washed with binding buffer three times before intravenous injection. Afterwards, a tail vein catheter containing 100 IE/ml heparin and 10 μg/ml mouse VEGF-A in sterile saline was inserted after anesthesia onset and prior to imaging. 10 μl heparin was directly injected, while 100 μl/30 g VEGF-A was injected after 1 min of baseline flow recording. Fast *xyt* imaging was performed with resonance scanner for every islet graft (Ex.: 561 & 633 nm, Em.: 558-564 nm & 650-750 nm). Around 20 RBCs were randomly picked at each time point for each islet, and individual RBC flow velocity was calculated by comparing two consecutive images.

### Image analysis

Unprocessed original images were used in the quantification of islet morphology and blood flow rate. Graft volume was estimated from backscatter signals, and dextran-labelled islet capillary structures were used for the estimation of vascular volume and diameters (Volocity, Perkin Elmer, USA), with same thresholds used for all groups at all time points. Relative islet volume of each graft in percentage was calculated from the values measured at individual time points and divided by the value at week 0. Relative vascular volume in percentage was calculated by dividing the volume of islet vessels by the estimated islet volume at each time point.

Time lapse confocal image stacks were analyzed and fluorescence intensity was quantified using Fiji[39] with the plugin Time Series Analyzer V3. For Ca^2+^ imaging, movements were corrected with a custom MATLAB program (Mathworks, USA), and all visible vessel segments were hand-picked for each islet. Dextran leakage measurement was performed as described before[23]. Islet borders were identified by backscatter signals, and average fluorescence intensity was measured in a ring structure of 30 pixels thickness outside each islet. The simulation of *in vivo* leakage kinetics was done in GraphPad Prism 9 using the model of plateau followed by one phase association.

### VEGF-A and sFlt-1 measurement

For measuring ex vivo VEGF-A production, isolated islets were cultured overnight. 30 islets from each group were picked into 4-well plates containing 450 μl culture medium per well. 72 hours later, supernatants were collected, briefly centrifuged and stored at −80 °C before measurement (Mouse VEGF Quantikine ELISA Kit, R&D Systems, USA). Islets were collected and lysed in 50 μl T-PER buffer (Thermo Fisher Scientific, USA) to quantify the DNA contents for normalization (Quant-iT PicoGreen dsDNA Assay Kit, Thermo Fisher Scientific, USA). 15 μl blood was taken from the tail vein from non-fasted animals around 9 A.M. for the measurement of plasma VEGF-A and sFlt-1(Mouse VEGFR1/Flt-1 Quantikine ELISA Kit, R&D Systems, USA) levels. Secreted VEGF-A in supernatants, plasma VEGF-A, sFlt-1 and islet DNA content were all measured according to manufacturers’ instructions.

### VEGFR-2 internalization

5×10^4^ HUVECs were plated on coverslips in 6 well plates and treated with 0.075% BSA and 25 mM Mannitol (BSA/Man) in the control group, or 50 μM palmitate (conjugated to BSA at a ratio of 4:1) and 25 mM D-glucose (Pal/D-glu) in the diet-mimicking group for 5 days. The recovery group was treated with Pal/D-glu for 2 days and BSA/Man for another 3 days. On day 6, confluent cells were starved for 5 hours with MV2 supplemented with 0.2% FBS. Starved cells were gently washed twice with PBS, and half of the cells were stimulated with 50 ng/ml human VEGF-A (Peprotech, Sweden) for 20 min at 37°C. VEGF-A containing medium was quickly removed afterwards and cells were washed twice in ice-cold PBS. Subsequently, cells were fixed in 4% paraformaldehyde solution at room temperature for 15 min and blocked with 2% BSA in PBS for 1 hour without permeabilization. Further incubations with anti-VEGFR2 N-terminal antibody (rabbit, 1:100, R&D Systems, USA) and goat anti-rabbit IgG H+L Alexa Fluor 488 (1:1000, Thermo Fisher Scientific, USA) were carried out at room temperature for 1 hour each, and cells were washed 3 times with PBS+/+ in between.

### Pancreatic sections and immunofluorescence staining

For staining of pERK1/2, animals were anesthetized with a mixture of 0.5 mg/kg fentanyl, 16 mg/kg fluanisone (Hypnorm, Roche, Switzerland) and 8 mg/kg midazolam (Hameln, Sweden), and were kept on a heating pad (37 °C) with oxygen supply throughout the experiment at 250 ml/min for 10 min. Afterwards, 100 μl/30 g mouse VEGF-A was injected through the tail vein and animals were kept on the heating pad for another 5 min before being sacrificed quickly by cervical dislocation. Pancreata were swiftly removed from euthanized animals and rinsed in cold PBS+/+, which took an additional minute before fixation by 4% paraformaldehyde in PBS+/+ at room temperature for 2 hours. Tissues were then washed with PBS+/+ twice and cryopreserved stepwise in 10%, 20% and eventually 30% sucrose in PBS+/+ at 4 °C overnight. They were embedded in O.C.T. freezing medium (Thermo Fisher Scientific, USA) at −80 °C and subsequently cut into 20 μm thick sections for staining. Sections were incubated with primary antibodies, including anti-VEGF-A antibody (goat, 1:50, R&D Systems, USA), anti-insulin antibody (rabbit, 1:500, Abcam, UK), anti-PECAM-1 antibody (goat, 1:400, R&D Systems, USA), anti-VEGFR2 (rabbit, 1:200, Cell Signaling Technology, USA), and phospho-p44/42 MAPK Thr202/Tyr204 antibody (rabbit, 1:200, Cell Signaling Technology, USA), at 4 °C overnight. After washing with PBS+/+ 3 times, samples were further incubated with secondary antibodies (donkey anti-rabbit IgG H+L Alexa Fluor 546 and donkey anti-goat IgG H+L Alexa Fluor 488, both at 1:1000, Thermo Fisher Scientific, USA) at 4 °C overnight and mounted with fluorescence mounting medium (Agilent, USA).

### Western Blots

Confluent HUVECs were stimulated by 50 ng/ml human VEGF-A for indicated lengths of time and lysed in M-PER buffer (Thermo Fisher Scientific, USA) supplemented with protease and phosphatase inhibitors (Mini Complete/PhosphoStop, Roche Diagnostics, Germany). Protein contents were determined using the BCA Protein Assay Kit (Pierce, USA). Protein extracts were separated on 7.5% stain-free precast gels (Bio-Rad, USA) and transferred to 0.22 μm PVDF membranes (GE Healthcare, USA). Blots were probed with primary antibodies, including phospho-VEGFR2 Y1175 antibody (rabbit, 1:1000), VEGFR2 antibody (rabbit, 1:1000), Phospho-PKCζ/λ (Thr410/403) antibody (rabbit, 1:500), PKCι/λ antibody (rabbit, 1:1000), phospho-Akt Thr308/473 antibody (rabbit, 1:1000), Akt1/2 antibody (rabbit, 1:1000), phospho-p44/42 MAPK Thr202/Tyr204 antibody (rabbit, 1:2000), Erk1/2 antibody (rabbit, 1:2000; all from Cell Signaling Technology, USA), and α-tubulin antibody (mouse,1:1000, Sigma-Aldrich, USA), and subsequently probed with secondary anti-mouse/-rabbit IgG (H+L)-HRP conjugates (1:6000, GE Healthcare, USA). Quantification was carried out on original scanned images using Image Lab software (Bio-Rad, USA) and normalization to total protein amount.

### In vitro perifusion of islets

Freshly isolated islets were kept in culture for 2 hours, and around 60 islets from each group were put in an automated perifusion system (BioRep Perifusion System, BioRep, USA) for the evaluation of dynamic insulin release *in vitro*, as described before [40]. Briefly, islets were immobilized in a gel (Bio-Gel P-4, Bio-Rad, USA) in flow chambers, and were perifused with specified solutions at a speed of 50 μl/min for indicated periods. Perfusates were continuously and automatically collected into 96-well plates for the detection of insulin (Insulin AlphaLISA Detection Kit, Perkin Elmer, USA). Insulin measurement was performed according to the manufacturer’s instructions.

### Transmission electron microscopy (TEM)

Islet grafts were dissected out from the eyes of euthanized recipient animals, and peri-islet iris tissues were removed carefully with micro scissors. Grafts were then fixed in 2.5% glutaraldehyde and 1% paraformaldehyde in 0.1 M phosphate buffer (pH 7.4) at 4 °C overnight. Samples were further washed in 0.1 M phosphate buffer (pH 7.4) and post-fixed in 2% osmium tetroxide 0.1 M phosphate buffer (pH 7.4) at 4 °C for 2 hours, dehydrated in ethanol and acetone, and embedded in LX-112 (Ladd Research Industries, Burlington, USA). Ultrathin sections of 50–60 nm thickness were made by Leica EM UC6 ultramicrotome (Leica Microsystems, Germany) and contrasted with uranyl acetate, followed by lead citrate treatment and examined in Tecnai 12 Spirit BioTWIN transmission electron microscope (FEI company, USA) at 100 kV. Images were acquired by 2kx2k Veleta OSiS CCD camera (Olympus Soft Imaging Solutions, Germany).

### Quantitative RT-PCR

Total RNA samples were extracted from freshly isolated islets with GeneJET RNA purification kit (Thermo Fisher Scientific, USA), and extracted mRNA was transcribed into cDNA using the Maxima First Strand cDNA Synthesis Kit (Thermo Fisher Scientific, USA), all according to manufacturer’s instructions. *Tbp*, *Vegfa, Kdr, Pecam1, L19, KDR, EFNB2, PARD3* and *DAB2* messages were measured using PowerUp SYBR Green Master Mix (Thermo Fisher Scientific, USA) in QuantStudio™ 5 Real-Time PCR System (Applied Biosystems, USA). Relative gene expression levels were quantified against the housekeep genes *Tbp* or *L19* messages and normalized to those of control islets or HUVECs respectively. All primer sequences are listed in Supplemental Table 1.

### Statistical tests

All results are presented as mean ± SEM (with individual data points for bar graphs). Conventional Two-way ANOVA or Two-way repeated measures ANOVA tests were used for time course analysis where appropriate. One-way ANOVA tests were used for comparison of multiple groups. Unpaired *t*-tests or Mann-Whitney tests were used for two group comparisons where appropriate, with a *p* value < 0.05 considered to indicate significance (GraphPad Prism 9).

## Results

### Western diet leads to body weight gain, glucose intolerance and pancreatic islet vascular remodeling

In order to depict the temporal dynamics of pancreatic islet vascular damage in a mouse model undergoing diet-induced obesity, we fed male C57BL/6J mice with either control diet (CD) or western diet (WD) and monitored obesity-related alterations in islet vascular morphology and functionality over time. We first transplanted 6-8-week-old mice with islets derived from wild type (WT) donors into their anterior chamber of the eye (ACE), and allocated recipients randomly into CD and WD groups after another 8 weeks, when islet engraftment and re-vascularization was complete (Figure 1A). The engrafted islets allow us to non-invasively and longitudinally investigate islet vascular biology at cellular level in vivo, as their newly formed vessels largely mirror characteristics of in situ islet vasculature[41, 42]. The diet intervention lasted for a total of 48 weeks, and recipient mice in both groups were examined every 4 weeks (Figure 1A). As expected, WD-fed mice rapidly developed truncal obesity, and showed slowing but continuous weight gain throughout the intervention, eventually gaining significantly more weight than the CD-fed mice from week 0 to the end point (Figure 1B and 1C). Meanwhile, glucose tolerance also deteriorated rapidly in WD group, although the level of intolerance stabilized after 8 weeks (Figure 1D and 1E), verifying the occurrence of metabolic disorders. We examined concurrently the morphological modifications of individual islet grafts and their vasculature over the course of diet intervention. Through intravenous injection of fluorescein isothiocyanate (FITC)–labelled dextran, we were able to visualize the reformed vasculature in grafted islets, and backscatter signals were obtained simultaneously for the estimation of islet size[23] (Figure 1F). Quantitative analysis showed obvious islet hyperplasia in the WD group (Figure 1G), accompanied by remarkable vascular growth and remodeling. The vessels of islet grafts in WD-fed mice displayed irregularly enlarged diameters starting from 12 weeks after diet intervention (Figure 1H), and they constituted a higher percentage of total islet volume (Figure 1I). By contrast, islet vasculature in CD group remained relatively stable, with limited growth and structural refinements (Figure 1F, 1H and 1I). Therefore, islet vessels are sensitive to diet manipulations and undergo continuous remodeling under WD intervention.

**Figure. 1.**
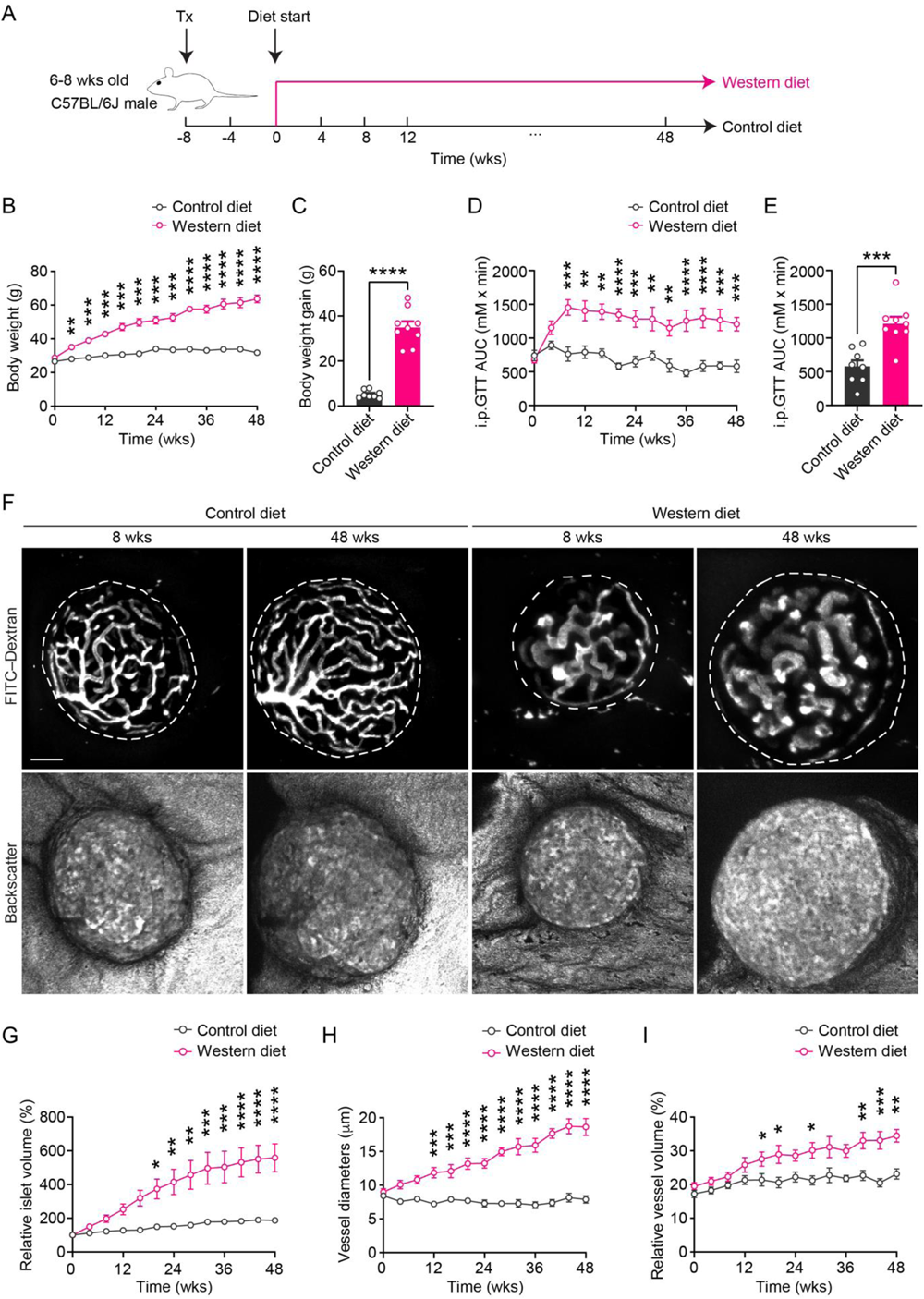
Western diet leads to body weight gain, glucose intolerance, islet hyperplasia and islet vascular remodeling. **A,** Schematic of experimental timeline. **B**-**C,** Body weight changes during diet intervention (B) and average body weight gain in CD- (n=8) and WD-fed (n=9) animals at week 48 (C). **D**-**E,** Area under the curve for intraperitoneal glucose tolerance test (i.p.GTT) during diet intervention (D) and at week 48 (E) in CD- (n=8) and WD-fed (n=9) animals. **F,** Morphological changes in islet grafts and vasculature over time in CD- and WD-fed animals. Representative confocal images are presented as maximum intensity projections. Scale bar: 50 μm. **G,** Relative growth of islet grafts in CD- (n=6) and WD-fed (n=7) animals during diet intervention compared to week 0. **H**-**I**, Vessel diameters (H) and relative vascular volume (I) in islet grafts of CD- (n=7) and WD-fed (n=9) animals during diet intervention. Data are shown as mean ± SEM (B, D, G, H and I) or individual points (C and E). Statistics are based on unpaired, two-tailed Student’s t tests (C and E) or two-way ANOVA (B, D, G, H and I), \**p*<0.05, \*\**p*<0.01, \*\*\**p*<0.001, \*\*\*\**p*<0.0001.

### Western diet enhances islet expression and production of VEGF-A

Vascular remodeling is usually accompanied by functional adaptations. To evaluate islet vascular functionality under WD feeding, we first examined islet expression of VEGF-A and of its receptor VEGFR2, which conveys the predominant signal that maintains the morphological and functional architecture of islet vessels[14, 16, 43]. Staining of pancreatic cryosections with antibodies against VEGF-A and VEGFR2 exhibited similar patterns of protein expression between CD- and WD-fed mice after 8 weeks of diet intervention (Figure S1A and S1B). Consistent with previous studies, VEGF-A was expressed by all islet endocrine cells, including non-β cells, at a much higher level than the surrounding exocrine cells in both groups (Figure S1A). VEGFR2 expression was mostly restricted to the membrane of islet vascular endothelial cells, as identified by the endothelial cell marker PECAM-1 (Figure S1B). With VEGF-A ligand produced by islet endocrine cells, they form an intra-islet paracrine signaling loop. Of note, VEGFR2 was also expressed at markedly lower level in exocrine pancreas (Figure S1B), implying that intra-islet endothelial cells are likely more sensitive than exocrine endothelial cells to perturbations in VEGF-A/VEGFR2 signaling under pathological conditions. Gene expression analysis showed that freshly isolated islets from WD group expressed more *Vegfa* gene after 12 weeks of diet intervention (Figure S1C). As a result, they released more VEGF-A ex vivo than the CD group (Figure S1D). The *Kdr* gene (encoding VEGFR2) was also upregulated in the WD group after 12 weeks of diet intervention compared to the CD group (Figure S1E). This may be attributed to a relatively augmented number of intra-islet endothelial cells, which is reflected by elevated *Pecam1* expression at the same time points (Figure S1F) and is consistent with our observation of increased vascular volume (Figure 1I). These changes in gene expression indicate that intra-islet VEGF-A/VEGFR2 paracrine signaling may be perturbed by WD intervention.

### Western diet diminishes VEGF-A triggered islet endothelial cell Ca^2+^ mobilization in vivo

One of the signaling events triggered by the binding of VEGF-A ligand to VEGFR2 is rapid mobilization of intracellular Ca^2+^, which serves as a suitable readout for VEGF-A/VEGFR2 signaling activity[44, 45]. Therefore, we generated a mouse line expressing a fluorescent reporter for intracellular Ca^2+^ concentration ([Ca^2+^]_i_) in endothelial cells. It was derived from the Ai38 line that carries a *loxP* flanked STOP cassette preventing the expression of a CAG promoter–driven Ca^2+^ biosensor GCaMP3 (GC3) in *Rosa26* locus[46]. Crossing this line to an endothelial cell–specific transgenic *Cdh5-Cre* mouse line led to the labelling of all VE-Cadherin–expressing endothelial cells with GC3 (EC-GC3) (Figure S2A)[47]. Intraperitoneal glucose tolerance tests on EC-GC3 animals showed no difference as compared to wild type animals in terms of glucose handling (Figure S2B). Immunostaining of pancreatic sections from EC-GC3 animals confirmed that GC3 primarily labelled the entire intra-islet vascular network, which was demonstrated by co-staining of PECAM-1 and GC3 (Figure S2C). When EC-GC3 animals were used as recipients for transplantation, islet grafts in the ACE of these mice recruited the GC3-expressing endothelial cells during re-vascularization (Figure S2D), making it an ideal model for monitoring the dynamics of [Ca^2+^]_i_ triggered by VEGF-A in vivo.

We then followed the previous timeline of investigation and divided EC-GC3 recipients into CD and WD groups 8 weeks after transplantation, when re-vascularization of engrafted reporter islets was complete[41] (Figure 2A). VEGF-A–induced [Ca^2+^]_i_ dynamics in islet endothelial cells was then assessed every 4 weeks until the end point to obtain a temporal overview of intra-islet VEGF-A/VEGFR2 signaling profile in the CD and WD groups respectively, and capture the emergence of any signaling disturbance. Baseline GC3 fluorescence in islet endothelial cells was generally low prior to stimulation and remained stable and of similar intensity in the CD and WD groups over time (Figure S3). When VEGF-A was administered intravenously, there was an instantaneous increase in GC3 fluorescence which peaked at about 1.5 minutes after injection and remained elevated for another 4 minutes before returning to baseline (Figure 2B). To quantitatively analyze this response, all visible intra-islet vascular segments were selected, and each [Ca^2+^]_i_ trace was plotted individually as fold increase over baseline in GC3 fluorescence intensity (Figure 2C). After 12 weeks of diet intervention, vessel segments within the same islet in the CD group typically had relatively uniform responses to VEGF-A, in the form of a distinct single raise (Figure 2D, left panel, Supplemental Movie 1). However, the vessel segments in the WD group showed diverse responses to VEGF-A, although no delay in the response was detected (Figure 2D, right panel, Supplemental Movie 2). While a few vessel segments in this group exhibited similar traces to the control segments, others had smaller peaks and/or shorter duration of response. A closer inspection of the peak responses from all animals in each group at 12 weeks of diet intervention revealed a distinct subgroup of vessel segments which were less responsive to VEGF-A in the WD-fed mice (Figure 2E). Consequently, when we averaged all [Ca^2+^]_i_ traces from individual vessel segments during the course of diet intervention, Ca^2+^ mobilization in the WD group became visibly smaller than that in the CD group at 12 weeks, and the difference grew more pronounced by the end time point (Figure 2F). Using area under the curve as a measure of total Ca^2+^ response, we observed a modest ageing-related decline in the CD-fed mice towards the end point (Figure 2G). By contrast, the WD group displayed a clear downward trend throughout the diet intervention, although there were signs of adaptation until week 32 as evidenced by the small fluctuations within the curve (Figure 2G). We thereby conclude that prolonged WD feeding diminishes VEGF-A–initiated signal transduction in islet endothelial cells, which is reflected by blunted Ca^2+^ response to a single VEGF-A bolus in vivo.

**Figure 2.**
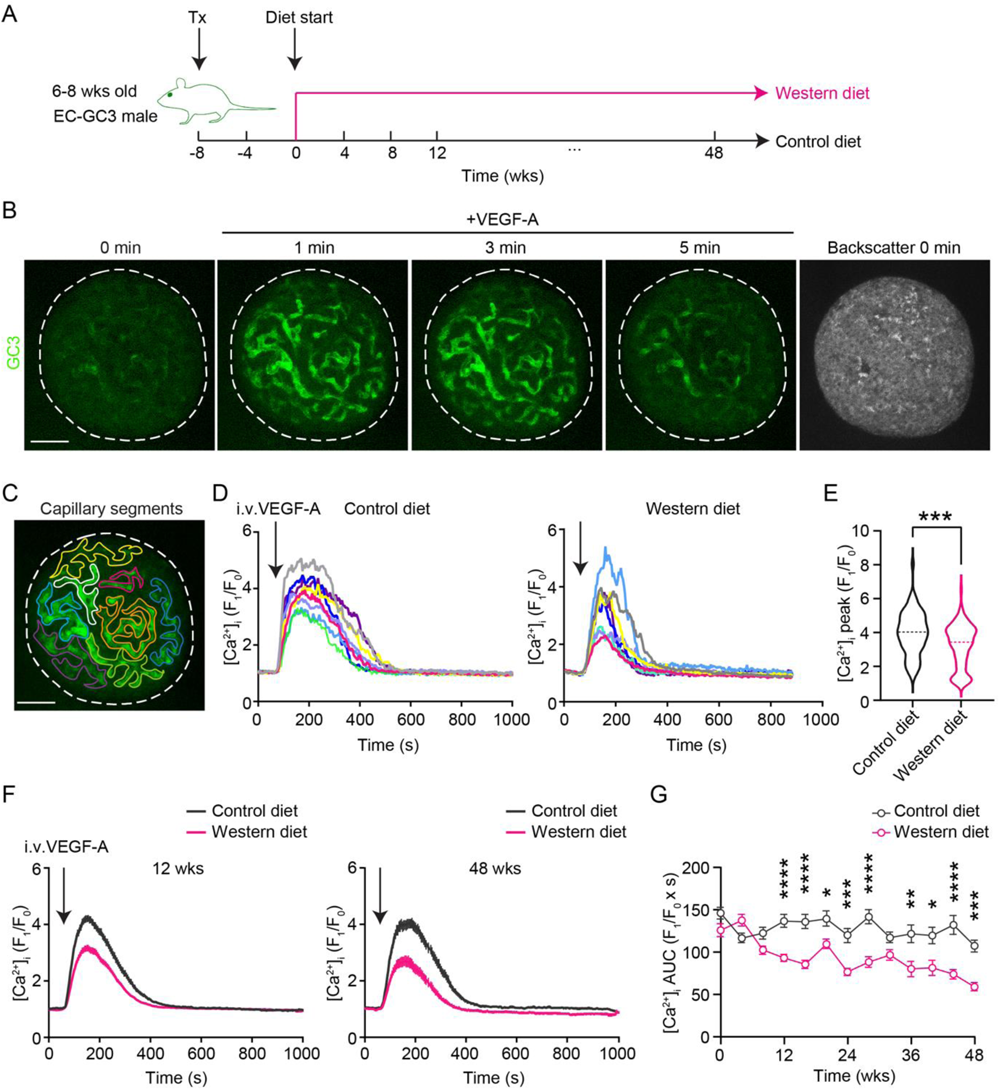
Western diet diminishes VEGF-A–triggered islet endothelial cell Ca^2+^ mobilization in vivo. **A,** Schematic of the experimental timeline. **B,** Representative intra-islet vessel Ca^2+^ response to VEGF-A bolus in the CD-fed animals. Scale bar: 50 μm. **C,** An example of vessel segment selection within an islet graft. Scale bar: 50 μm. **D**, Representative Ca^2+^ traces from individual segments within a single islet in CD- (left) and WD-fed (right) animals at week 12. **E,** Distribution of peak Ca^2+^ response in CD- (n=8) and WD-fed (n=12) animals at week 12. **F**, Averaged Ca^2+^ traces in CD- and WD-fed animals at week 12 (left, CD: n=8; WD: n=12) and week 48 (right, CD: n=6; WD: n=6). **G**, Area under the curve for Ca^2+^ traces in CD- (n=6-12) and WD-fed (n=6-13) animals during diet intervention. Data are shown as violin plot (E) or mean ± SEM (the rest). Statistics are based on Mann-Whitney test (E) or two-way ANOVA (G), \**p*<0.05, \*\**p*<0.01, \*\*\**p*<0.001, \*\*\*\**p*<0.0001.

### Western diet undermines VEGF-A regulation of islet hemodynamics

VEGF-A–triggered intracellular Ca^2+^ mobilization further elicits endothelium-dependent vasodilation through rapid generation of nitric oxide[48–50], which is counteracted by increased endothelin-1 release and consequent vasoconstriction[51–53]. Together with neuronal and nutritional factors, the balance of these complex signaling cascades shapes the pattern of highly dynamic islet perfusion, which is normally coupled to the metabolic demand and crucial for timely glycaemic control[54–57]. In this respect, we investigated if the vasoactive effect of VEGF-A could also be undermined by WD feeding. We used red blood cell (RBC) flux as a direct measure for the speed of islet blood flow and examined the real-time fluctuations brought by VEGF-A application in vivo. A small number of RBCs in each mouse was pre-labelled by a lipophilic fluorescent dye[38], and the movement of individual RBCs was traced in fast time-series scans for estimating the velocity of local blood flow (Figure 3A and Supplemental Movie 3). Baseline blood flow velocity was moderately higher in WD-fed mice after 12 weeks of feeding (Figure 3B), which may be due to elevated systemic blood pressure[58]. An instant decrease in islet RBC flow velocity upon intravenous VEGF-A injection was evident in CD-fed mice, which reached its minimum 2 min later and slowly recovered to baseline level approximately 10 min after the injection (Figure 3C). By contrast, islet blood flow in WD-fed mice at 12 weeks was not as sensitive to VEGF-A manipulation as compared to control, and the sharp dip in flow velocity was attenuated (Figure 3C and 3D). Taken together, disrupted VEGF-A regulation of islet hemodynamics verifies that islet endothelial cells are rendered less responsive to VEGF-A in WD-fed mice, implying that the fine regulation of islet blood flow may be compromised.

**Figure 3.**
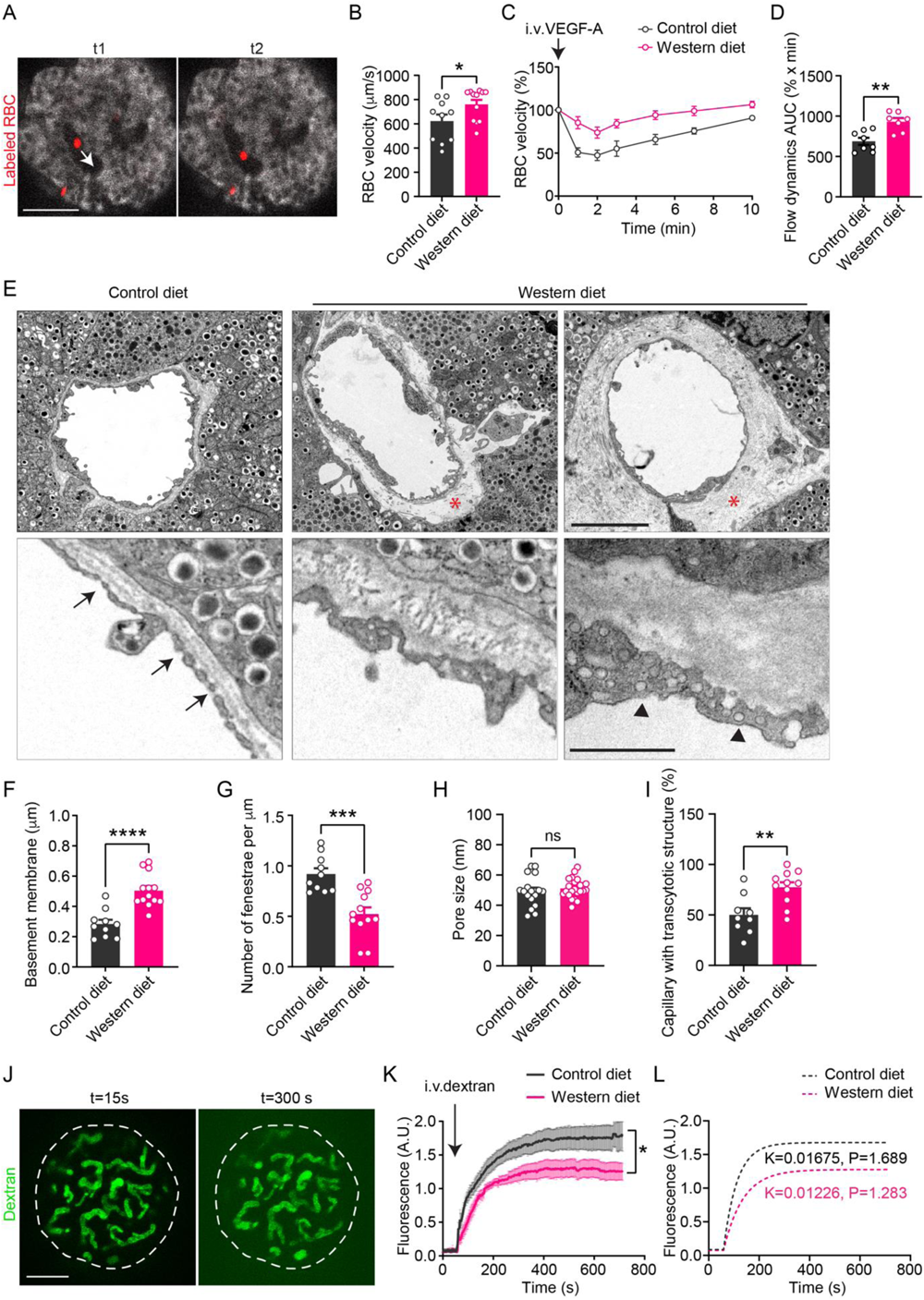
Western diet undermines VEGF-A regulation of islet hemodynamics and islet vascular barrier function. **A**, Movement tracking of labelled individual RBCs (red) in the blood stream in two consecutive time-series images (time interval between t1 and t2 is 18 ms). Arrow indicates the direction of blood flow. Scale bar: 50 μm. **B**, Baseline RBC velocity in CD- (n=10) and WD-fed (n=12) animals at week 12. **C,** Average RBC velocity dynamics upon VEGF-A injection in CD- (n=8) and WD-fed (n=7) animals at week 12. **D**, Area under the curve for RBC velocity excursions in CD- (n=8) and WD-fed (n=7) animals at week 12. **E**, Electron micrographs of islet grafts dissected from the ACE of CD- and WD-fed animals at week 12. Scale bars: 5 μm (upper) and 1 μm (lower). **F-I**, Basement membrane thickness (F), number of fenestrae (G), pore sizes of fenestrae (H) and percentage of capillaries with transcytotic structure (I) in islet vessels of CD- (n=9) and WD-fed (n=11) animals at week 12. **J,** Representative images showing distribution of FITC-dextran inside and around islet grafts 15s (left) and 300s (right) after intravenous injection. Scale bar: 50 μm. **K**-**L**, Average dextran fluorescence intensity dynamics outside islet grafts upon injection in CD- (n=12) and WD-fed (n=10) animals at week 12 (K) and simulated curves showing different kinetics of leakage (L). Data are shown as mean ± SEM (C and K) or individual points (B, D, F, G, H and I). Statistics are based on unpaired, two-tailed Student’s t tests (B, D, F, G, H and I) or two-way ANOVA (K), \**p*<0.05, \*\**p*<0.01, \*\*\**p*<0.001, \*\*\*\**p*<0.0001 and ns (\**p*>0.05).

### Western diet causes ultrastructural changes in islet vessels and alters vascular permeability

Apart from the dynamic islet blood flow, another vascular element that affects the efficiency of islet action is the barrier function, which largely relies on the permeability of islet endothelial cells and the unique ultrastructure of islet vessels, both being under the regulation of VEGF-A[59–61]. In mouse islets, endocrine cells are separated from vascular cells by a layer of basement membrane, the deposition of which is modulated by VEGF-A[19, 62]. Also, the islet vessels have 10 times more diaphragmed fenestrae than those in the exocrine pancreas, which are primarily maintained by active VEGF-A signaling[14, 16, 63]. To study these properties, we dissected out islet grafts from the ACE of CD- and WD-fed mice at given time points and performed transmission electron microscopy analysis on the ultrastructure of islet vessels. Images derived from the CD group showed a nice thin layer of basement membrane between endothelial cells and granulated endocrine cells (Figure 3E, upper left panel). However, this layer was significantly and often unevenly thickened in the WD group at 12 weeks (Figure 3E, upper middle and right panels; Figure 3F), which not only increased the travel distance for molecules between islet endocrine cells and the blood stream, but also added to the stiffness of islet vascular walls, undermining their elasticity[64]. The endothelial cell lining of islet vessels was densely fenestrated in control islet grafts (Figure 3E, lower left panel), whereas the number of fenestrae was notably smaller in the WD group (Figure 3E, lower middle panel; Figure 3G), although the pore sizes in both groups were comparable (Figure 3H, approximately 50 nm on average). In addition, we noticed more caveolae-like transcytotic structures in islet vessels from the WD group compared to the CD group (Figure 3E, lower right panel; Figure 3I), probably serving as compensation for the loss of fenestration.

The fenestrated islet endothelium is highly permeable and allows bidirectional transfer of solutes and molecules such as small peptides through the pores along the endothelial cell lining. This feature enables quick equilibration between the islet interstitial fluid and blood stream, thus facilitating islet sensing of glucose as well as insulin disposal. Considering the ultrastructural alterations and the VEGF-A signaling blockage which could hamper its permeabilizing effect, we investigated if the islet vessel permeability and barrier function in the WD-fed mice were compromised. As previously described[23], we gave mice in the CD and WD groups a single intravenous injection of 40 kDa FITC-labelled dextran and measured its diffusion into the aqueous humour outside the islet graft over time (Figure 3J). We chose this as a measure for islet vessel permeability, since 40 kDa dextran preferentially leaks through fenestrated islet vessels instead of the less permeable iris vessels. Upon injection of dextran, there was a rapid increase of FITC fluorescence outside the islet grafts, followed by a plateau that was established approximately 6 minutes later, as a result of the active drainage of aqueous humour, renal excretion and a declining dextran concentration gradient[65, 66]. Although both groups reached the plateau at matching speeds, the initial rate of leakage and the final height of the plateau were significantly reduced in islet grafts from the WD-fed mice (Figure 3K). Simulation of this process as previously described[23] also generated two distinct curves, which supports different kinetics of dextran leakage in CD and WD groups (Figure 3L). Altogether, these results suggest that islet vessels in WD-fed mice are markedly less permeable to medium sized molecules, which could potentially pose an additional obstacle for secreted insulin to reach the blood stream.

### Dysfunctional islet vessels hinder insulin outflow in vivo despite normal secretory capacity of islet β cells

To determine the actual effects of impaired islet vascular barrier function on glucose homeostasis, we compared insulin secretion and distribution kinetics of islets from CD- and WD-fed mice. Freshly isolated islets from mice fed with CD or WD for 12 weeks were cultivated for 2 hours before being packed in columns that were connected to an in vitro perifusion system. The shortened cultivation period minimizes changes in β cell gene and protein expression, so that the glucose sensing and insulin secretory machinery resembles in situ condition as closely as possible. Packed islets were subsequently perifused with different concentrations of glucose as well as KCl, and the perfusate was continuously collected. In this case, solutions containing secretagogues reach β cells via diffusion, and secreted insulin was released from islets directly into the perfusate, both without the need for islet vessels. Dynamic insulin secretion in response to 11 mM glucose was identical in islets from CD- and WD-fed mice (Figure 4A and 4B), suggesting that after 12 weeks of WD intervention, β cell response to glucose per se remains intact when they are separated from the unhealthy inner environment created by diet-induced obesity. We then examined insulin release kinetics in vivo during an intravenous glucose tolerance test, where well-controlled glucose delivery to β cells as well as insulin outflow into the blood stream rely on the native islet vasculature. As predicted, WD-fed animals displayed altered blood glucose excursion after 4 weeks of WD feeding (Figure 4C), and they stayed at similar levels of glucose intolerance during diet intervention (Figure 4D). After 12 weeks of WD feeding, mice had higher average fasting insulin levels than controls, whereas no delay in insulin secretion was observed (Figure 4E). Plasma insulin levels peaked at 1 min after glucose injection and dropped quickly after 3 minutes in both groups. Nevertheless, the average plasma insulin concentration at 1 minute was markedly lower in WD-fed mice despite higher baseline concentration (Figure 4E). Specifically, the elevation of circulating insulin concentration during the first minute of glucose injection was less prominent in WD-fed mice, in contrast to the sharper peak in CD-fed mice (Figure 4F). The amount of secreted insulin that crossed the endothelial barrier into the blood stream within the first 3 minutes of glucose challenge (measured as area under the curve) was also smaller in WD-fed mice (Figure 4G). Therefore, in spite of β cell functional resilience to diet intervention, the efficiency of islet insulin disposal was compromised by obstructed vascular routes in WD-fed mice, contributing to their dysregulated glucose handling.

**Figure 4.**
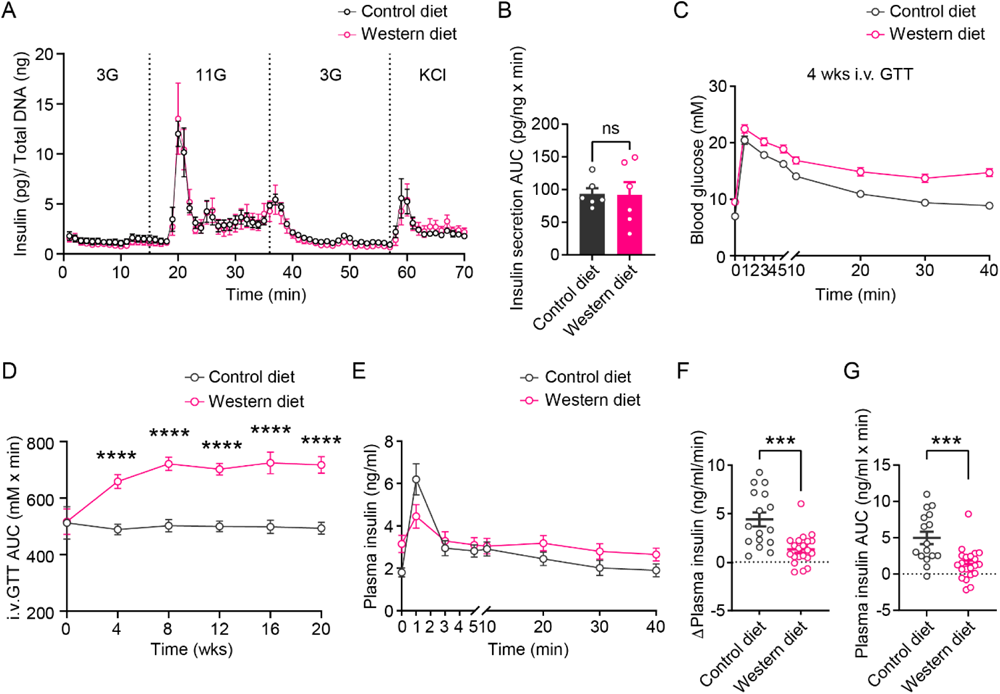
Dysfunctional islet vessels hinder insulin release in vivo. **A,** Dynamic insulin secretion from freshly isolated islets in a perifusion system. Islets were obtained from CD- and WD-fed animals at week 12 (n=3). **B,** Area under the curve for insulin secretion induced by 11 mM glucose (n=3). **C,** Intravenous glucose tolerance test (i.v.GTT) in CD- (n=18) and WD-fed (n=16) animals at week 4. **D,** Area under the curve for i.v.GTT in CD- (n=7-18) and WD-fed (n=6-23) animals from week 0 to week 20. **E**, Plasma insulin excursions during i.v.GTT in CD- (n=16) and WD-fed (n=21) animals at week 12. **F**, Increase in plasma insulin concentrations during the first min of i.v.GTT in CD- (n=16) and WD-fed (n=21) animals at week 12. **G,** Area under the curve for insulin excursions during the first 3 min of i.v.GTT in CD- (n=16) and WD-fed (n=21) animals at week 12. Data are shown as mean ± SEM (A, C, D and E) or individual points (B, F and G). Statistics are based on unpaired, two-tailed Student’s t test (B), two-way ANOVA (D) or Mann-Whitney tests (F and G), \**p*<0.05, \*\**p*<0.01, \*\*\**p*<0.001, \*\*\*\**p*<0.0001 and ns (\**p*>0.05).

### Islet pathogenesis elicited by WD is only partly reversible by diet switch

Islet endocrine cells exhibit a remarkable level of plasticity when exposed to enhanced metabolic stress. In particular, both β cell mass and functionality can be modulated to meet an increased need for insulin in obese and diabetic mouse models[67, 68]. Conversely, vascular endothelial cells are highly susceptible to glucotoxicity and lipotoxicity that are commonly present in obesity and diabetes[9, 10, 69]. To deepen our understanding of islet endothelial cell damage by WD and its possible reversibility, we added another diet group which was initially fed with WD for 24 weeks and then switched to CD for the remaining 24 weeks (Refed, Figure 5A). Mice in refed group quickly lost the excess body weight gained while they were on WD, and became as lean as control mice after 4 weeks (Figure 5B and 5C). Consequently, they displayed normal whole-body sensitivity to insulin (Figure S4), and intraperitoneal glucose tolerance was promptly improved (Figure 5D and 5E). Nonetheless, islet graft expansion throughout the entire time course remained more pronounced in refed mice than CD-fed mice, although the difference was not as striking as that between the CD and WD groups (Figure 5F and 5G). Furthermore, after 24 weeks of refeeding, the vasculature of islet grafts still resembled that of WD-fed mice (Figure 5F). The average vessel diameter in the refed group approximated that of the WD group and remained notably larger than the CD group (Figure 5H). Intriguingly, the relative vascular volume of refed group became indiscernible from that of the CD group and smaller than the WD group (Figure 5I), which may be due to decelerated growth of islet vessels in refed mice. Gene expression analysis on islets derived from all the three groups showed that *Vegfa* and *Kdr* expression was normalized by refeeding (Figure 5J and 5K), and islet VEGF-A production was also restored to that of the control level (Figure 5L). Altogether, visible improvements in whole-body metabolic status were noted in mice which were refed on CD, and pathological changes in islet *Vegfa* and *Kdr* gene expression elicited by WD were reversed. Nevertheless, rescue by diet switch appears to be insufficient for the full recovery of the damaged islet vasculature.

**Figure 5.**
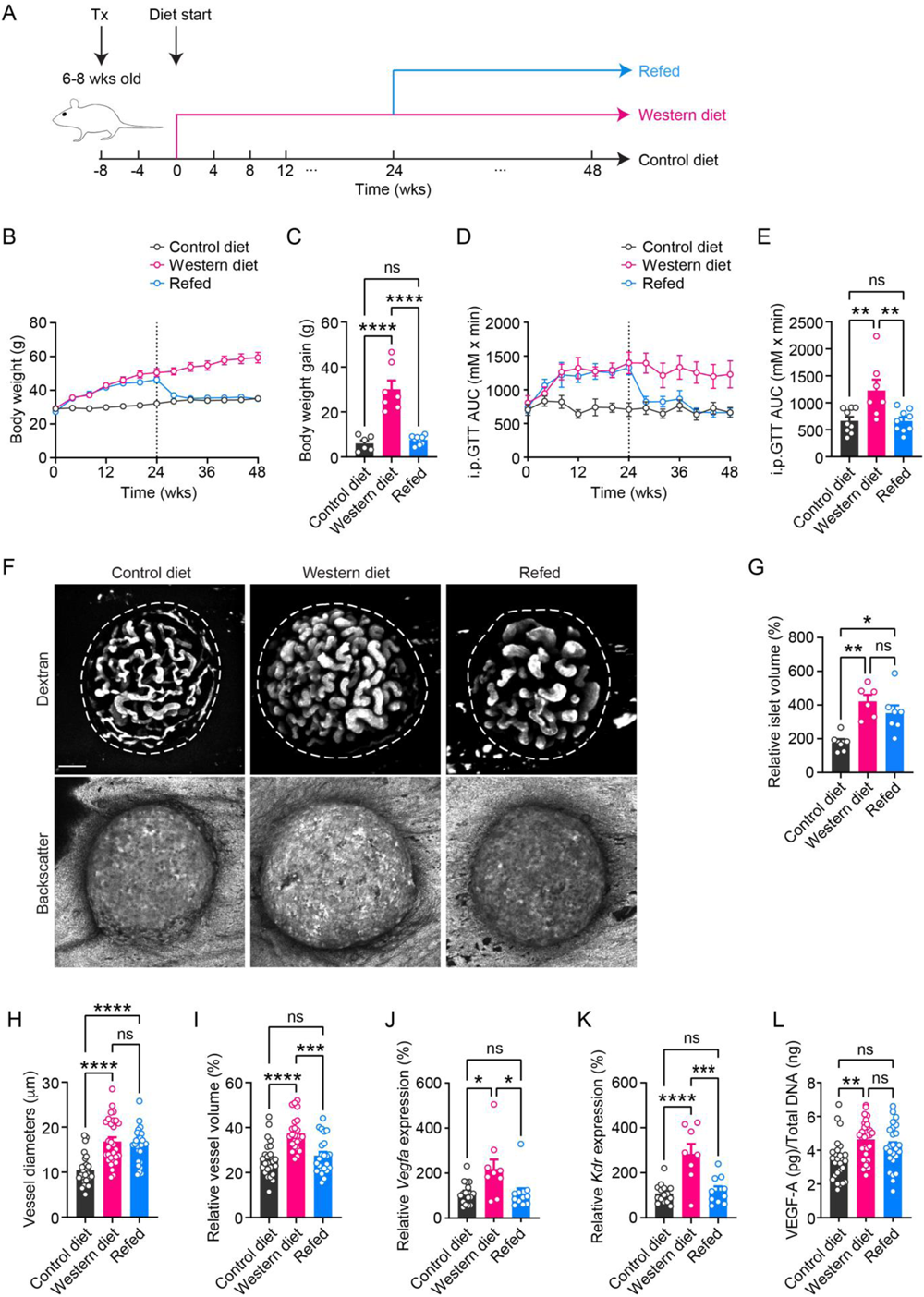
Diet reversal partly restores islet pathogenesis. **A,** Schematic of the experimental timeline. **B**-**C,** Body weight changes during diet intervention (B) and the average body weight gain in animals from CD (n=6), WD (n=7) and refed (n=9) groups at week 48 (C). **D**-**E**, Area under the curve for i.p GTT during diet intervention (D) and at week 48 (E) in animals from CD (n=9), WD (n=7) and refed (n=9) groups. **F,** Morphology of islet grafts and their vasculature in CD, WD and refed groups of animals at week 48. Representative confocal images are presented as maximum intensity projections. Scale bar: 50 μm. **G**, Relative growth of islet grafts in animals from CD (n=6), WD (n=6) and refed (n=7) groups at week 48 compared to week 0. **H**-**I**, Vessel diameters (H) and relative vascular volume (I) in islet grafts of animals from CD (n=7), WD (n=6) and refed (n=9) groups at week 48. **J**-**K,** Gene expression levels of *Vegfa* (J) and *Kdr* (K) in freshly isolated islets from CD (n=11-14), WD (n=8-9) and refed (n=10-11) groups of animals at week 48. **L,** VEGF-A production in cultured islets from CD (n=6), WD (n=6) and refed (n=8) groups of animals at week 48. Data are shown as mean ± SEM (B and D) or individual points (C, E, G, H, I, J, K and L). Statistics are based on one-way ANOVA (C, E, G, H, I, J, K and L), \**p*<0.05, \*\**p*<0.01, \*\*\**p*<0.001, \*\*\*\**p*<0.0001 and ns (\**p*>0.05).

### VEGF-A desensitization and ultrastructure-related barrier function impairment persist in islet endothelial cells after diet switch

In addition to vascular morphology, we re-evaluated all the parameters for islet vessel functionality after 24 weeks of refeeding. Surprisingly, VEGF-A–triggered Ca^2+^ mobilization in islet endothelial cells in the refed group of mice remained almost as blunted as in the WD group (Figure 6A). Both total Ca^2+^ response and the peak amplitude of [Ca^2+^]_i_ stayed considerably lower than in CD-fed mice (Figure 6B and 6C). Similarly, VEGF-A–induced hemodynamic changes remained diminished in the refed group, with no detectable improvement as compared to the WD group (Figure 6D and 6E). Jointly, there is no sign of islet endothelial cell recovery in terms of VEGF-A sensitivity by the end of refeeding. We then proceeded with the investigation of islet vessel ultrastructural and barrier properties at the same time point. Electron micrographs derived from micro-dissected islet grafts showed drastic thickening of basement membrane due to excessive deposition of matrix proteins in both WD and refed group of mice at 48 weeks (Figure 6F, upper panels; Figure 6G). Islet vessels in the refed group had notably more fenestration than WD-fed mice, although still less than the control group (Figure 6F, lower panels; Figure 6H), and the size of the pores were indiscernible among all three groups (Figure 6I). Moreover, the compensatory formation of transcytotic vesicles was reduced by refeeding, possibly due to an increased degree of fenestration (Figure 6J). Overall, with regard to the remaining differences between the control and refed group of mice, the islet vascular ultrastructure in the latter was not fully normalized. As a consequence, permeability to 40 kDa FITC-dextran in refed group was slightly higher but not significantly different from that of the WD group (Figure 6K), which was also supported by simulation (Figure 6L). Taken together, WD related desensitization to VEGF-A in islet endothelial cells was not simply reversed by diet switch. The ultrastructural abnormities sustained after 24 weeks of refeeding, indicating that islet vessels had not fully regained their functionality and the barrier function remained partially compromised. These results indicate that islet endothelial cells are indeed more vulnerable to WD-elicited toxicity than endocrine cells, while the inflicted damage is also long-lasting.

**Figure 6.**
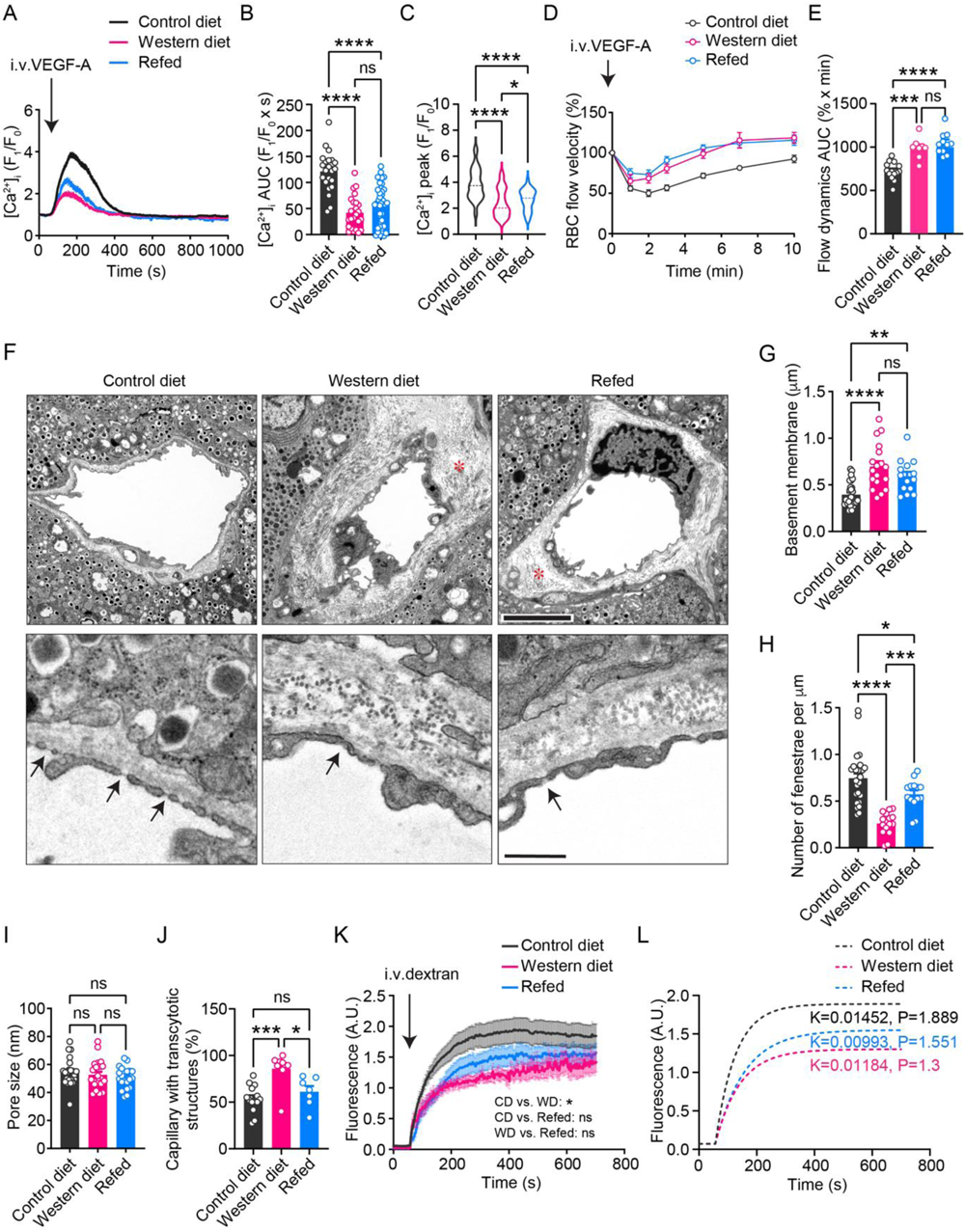
VEGF-A desensitization and ultrastructure-related barrier function impairment persist in islet vessels after refeeding. **A,** Averaged Ca^2+^ traces in animals from CD (n=4), WD (n=4) and refed (n=6) groups at week 48. **B,** Area under the curve for Ca^2+^ traces in animals from CD (n=4), WD (n=4) and refed (b=6) groups at week 48. **C,** Distribution of peak Ca^2+^ response in animals from CD (n=4), WD (n=4) and refed (n=6) groups at week 48. **D,** Average RBC velocity dynamics upon VEGFA injection in animals from CD (n=14), WD (n=8) and refed (n=10) groups at week 48. **E,** Area under the curve for RBC velocity excursions in animals from CD (n=14), WD (n=8) and refed (n=10) groups at week 48. **F,** Electron micrographs of islet grafts dissected from the ACE of CD, WD and refed groups of animals at week 48. Scale bars: 5 μm (upper) and 1 μm (lower). **G-J,** Basement membrane thickness (G), number of fenestrae (H), pore sizes of fenestrae (I) and percentage of capillaries with transcytotic structure (J) in islet vessels of CD (n=14), WD (n=9) and refed (n=7) groups of animals at week 48. **K**-**L,** Average dextran fluorescence intensity dynamics outside islet grafts upon injection in animals from CD (n=26), WD (n=14) and refed (n=13) groups at week 48 (K) and simulated curves showing different kinetics (L). Data are shown as mean ± SEM (A, D and K), individual points (B, E, G, H, I and J) or violin plot (C). Statistics are based on one-way ANOVA (B, C, E, G, H, I and J) or two-way ANOVA (K), \**p*<0.05, \*\**p*<0.01, \*\*\**p*<0.001, \*\*\*\**p*<0.0001 and ns (\**p*>0.05).

### Irreversible islet vessel dysfunction impedes insulin outflow despite preserved β cell secretory capacity

We have demonstrated that islet vessel dysfunction alone is detrimental to insulin output, aggravating glucose homeostasis in WD-fed mice. To further characterize the effect of dysfunctional islet vessels on glucose handling in refed mice, which fully recovered their body weight, intraperitoneal glucose tolerance as well as insulin sensitivity, we subjected freshly isolated islets from all three groups to the same in vitro perifusion system as described before for the investigation of dynamic insulin release under glucose challenge. At the end of diet intervention, the glucose stimulus-response curves were identical in islets derived from control and refed mice, while islets from the WD group showed slightly but not significantly reduced insulin secretion under glucose stimulation (Figure 7A and 7B). Therefore, islets from refed mice are capable of secreting as much insulin as control islets under the same glucose concentration, and β cell secretion is preserved by refeeding. However, intravenous glucose tolerance tests revealed a mildly impaired glucose handling in refed mice. They remained less efficient than control mice in glucose clearance, although exhibiting significant improvements from WD-fed mice (Figure 7C and 7D). This was consistent with blunted plasma insulin excursions in refed mice within the first 3 minutes of glucose injection, which resembled the WD group instead of the control (Figure 7E). The elevation of plasma insulin concentration at 1 minute was considerably lower in refed mice (Figure 7F), and the amount of insulin released into the blood stream within the first 3 minutes was also markedly less than control (Figure 7G). Meanwhile, both parameters in the refed group were comparable to the WD group. In summary, the irreversibly undermined islet vascular function alone poses a moderate but significant impediment to insulin release from islet β cells in vivo, thus hampering glucose clearance in the lean refed mice with intact islet function.

**Figure 7.**
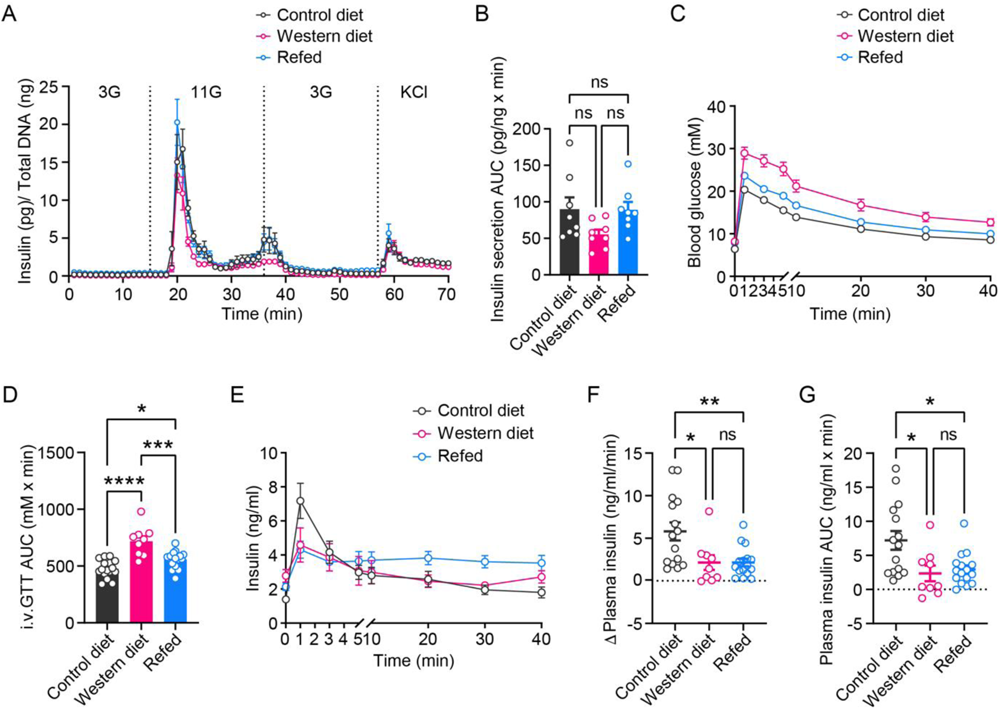
Irreversible islet vessel dysfunction impedes insulin release in vivo. **A,** Dynamic insulin secretion from freshly isolated islets in a perifusion system. Islets were derived from CD, WD and refed groups of animals at week 48 (n=4). **B,** Area under the curve for insulin secretion induced by 11 mM glucose (n=4). **C,** i.v.GTT in animals from CD (n=15), WD (n=9) and refed (n=16) groups at week 48. **D,** Area under the curve for i.v.GTT in CD (n=15), WD (n=9) and refed (n=16) groups of animals at week 48. **E,** Plasma insulin excursions during i.v.GTT in CD (n=15), WD (n=9) and refed (n=16) groups of animals at week 48. **F**, Increase in plasma insulin concentrations during the first min of i.v.GTT in CD (n=15), WD (n=9) and refed (n=16) groups of animals at week 48. **G,** Area under the curve for insulin excursions during the first 3 min of i.v.GTT in CD (n=15), WD (n=9) and refed (n=16) groups of animals at week 48. Data are shown as mean ± SEM (A, C and E) or individual points (B, D, F and G). Statistics are based on one-way ANOVA (B, D, F and G), \**p*<0.05, \*\**p*<0.01, \*\*\**p*<0.001, \*\*\*\**p*<0.0001 and ns (\**p*>0.05).

### Excessive glucose and free fatty acids diminish VEGF-A triggered VEGFR2 internalization and downstream signaling in endothelial cells

In order to gain an in-depth understanding of the mechanisms underlying islet vessel dysfunction caused by WD feeding, we examined several factors that may affect the strength of VEGF-A triggered signaling in islet endothelial cells. The soluble form of VEGF receptor 1 (sFlt-1) acts as a decoy VEGF-A receptor and binds the ligand at a much higher affinity than VEGFR2[70]. It is thus generally considered as a prominent antagonist to VEGF-A signaling. An elevated circulating level of sFlt-1 is linked to pathological situations such as preeclampsia[71], and was also observed in aging[72]. However, average plasma sFlt-1 concentration in mice fed with WD for 24 weeks didn’t differ from that of mice fed with CD (Figure S5A). Similarly, plasma sFlt-1 levels were also indistinguishable among CD, WD and refed groups of mice by the end point week 48 (Figure S5B). We can thus exclude the possibility that a lack of VEGF-A sensitivity in the WD and refed groups of mice is due to excessive neutralization of VEGF-A ligand by circulating sFlt-1. VEGFR2 internalization triggered by ligand binding is a crucial first step for full activation of its kinase activity[13]. This process relies on the association of VEGFR2 to the transmembrane Eph ligand ephrin-B2 via the clathrin-associated sorting protein Disabled 2 (Dab2) as well as the cell polarity protein PAR-3[73, 74], while the formation of this sorting complex is negatively regulated by aPKC through phosphorylation of Dab2[74]. We thus investigated the pattern of VEGFR2 activation and its downstream signaling cascades in cultured human endothelial cells. HUVECs were cultured in the presence of 50 μM palmitate and 25 mM D-glucose (Pal/D-glu) for 5 days to mimic overnutrition in WD-fed mice, while fatty acid–free BSA and Mannitol (BSA/Man) were used as control ingredients. In the “recovery” group, HUVECs were exposed to Pal/D-glu for 2 days followed by cultivation under control conditions for another 3 days. Immunofluorescence staining against the extracellular domain of VEGFR2 showed a diffuse distribution pattern over the cell membrane prior to stimulation in control HUVECs. After 20 min of incubation with VEGF-A, VEGFR2 fluorescence became more punctate and the total membrane fluorescence intensity decreased, due to receptor dimerization and internalization (Figure 8A, left panel; Figure 8B). However, in the Pal/D-glu and recovery groups, despite similar initial VEGFR2 fluorescence to that of the control cells, the decline in cell surface VEGFR2 fluorescence after VEGF-A stimulation was not as striking (Figure 8A, middle and right panels; Figure 8B), indicating reduced VEGFR2 internalization upon activation. We then investigated gene expression levels of *EFNB2*, *PARD3* and *DAB2* (encoding ephrin-B2, PAR-3 and Dab2 respectively) in HUVECs, and found no significant changes in any of these genes by Pal/D-glu treatment (Figure S5C). Meanwhile, control HUVECs have a low baseline level of activated aPKC, and its phosphorylation directed towards an increase upon VEGF-A stimulation (Figure 8C and 8D), consistent with previous reports[74]. Nevertheless, the baseline aPKC phosphorylation was significantly higher in HUVECs treated with Pal/D-glu (Figure 8C and 8D), which may diminish the formation of VEGFR2/Dab2/PAR-3/ephrin-B2 complex and thus hinder VEGFR2 internalization. HUVECs in the recovery group showed similar baseline aPKC phosphorylation level to that of the Pal/D-glu group (Figure 8C and 8D), implying that its hyperactivity was not restored by the removal of Pal/D-glu. Since activation of major VEGFR2 downstream molecules requires signal transmitted from the internalized receptors that are sorted into intracellular compartments such as early endosomes[13], we further compared VEGF-A induced phosphorylation of protein kinase B (AKT) and mitogen-activated protein kinase ERK1/2 under the above-mentioned experimental conditions (Figure 8E). Although phosphorylation of VEGFR2 at Tyr1175 didn’t seem to be affected by Pal/D-glu treatment (Figure S5D), activation of both AKT and ERK1/2 were markedly blunted in both the pal/D-glu and recovery groups, compared to control HUVECs (Figure 8E, 8F and 8G). Finally, to validate these findings in situ, pancreata from mice in CD, WD and refed groups at week 48 were harvested 6 minutes after an intravenous injection of VEGF-A and fixed immediately to capture the phosphorylation status of various signaling molecules downstream of VEGFR2. Immunofluorescence staining of pancreatic sections from CD-fed mice exhibited clear ERK1/2 phosphorylation and nuclear localization, while the number of pERK1/2 positive nuclei was visibly reduced, and the fluorescence signal was weaker in WD-fed and refed mice (Figure 8H). The percentage of pERK1/2 positive islet vessel area is thus highest in CD-fed mice, and indiscernible between WD and refed groups (Figure 8I). Overall, these results suggest that deficient VEGFR2 internalization due to over-activation of aPKC is an underlying mechanism for the irreversible islet vessel VEGF-A desensitization and dysfunction as a result of WD feeding.

**Figure 8.**
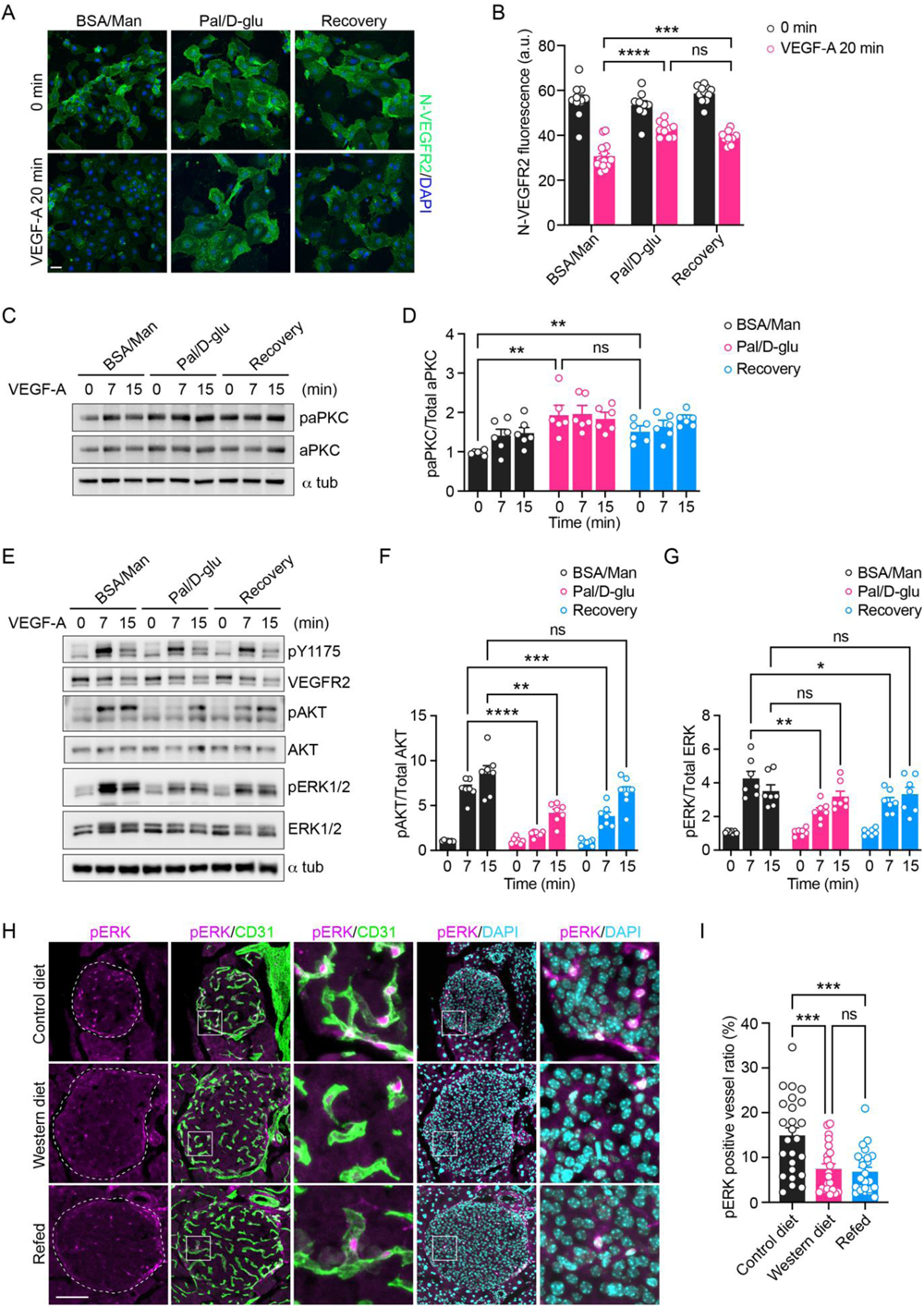
Western diet components impair VEGFR2 internalization and downstream signal transduction. **A,** Immunofluorescence staining of HUVECs cultivated under control, diabetic or recovery conditions, showing cell surface VEGFR2 (green) abundance before and 20 min after VEGF-A stimulation. Scale bar: 50 μm. **B,** Quantification of relative fluorescence intensity from cell surface VEGFR2 staining in HUVEC cultivated under control, diabetic or recovery conditions (n=3). **C,** Western blots showing phosphorylated and total aPKC levels in HUVECs cultivated under BSA/Man, Pal/D-glu or recovery conditions during VEGF-A stimulation. **D,** Normalized aPKC phosphorylation levels in HUVECs under control, diabetic or recovery conditions (n=3). **E,** Western blots showing VEGF-A induced phosphorylation of downstream signaling molecules in HUVECs cultivated under BSA/Man, Pal/D-glu or recovery conditions. **F-G,** Normalized AKT (F) and ERK1/2 (G) phosphorylation levels during VEGF-A stimulation in HUVECs cultivated under BSA/Man, Pal/D-glu or recovery conditions (n=4). **H,** Immunofluorescence staining of pancreatic sections from CD, WD and refed groups of animals at weeks 48, showing ERK1/2 phosphorylation (magenta) in islet vessels induced by intravenous VEGF-A injection. PECAM-1 and DAPI signals are shown in green and cyan respectively. Squares indicate areas magnified in the respective panels on the right side. Scale bar: 100 μm. **I,** Relative area of islet vessels that are positive for pERK1/2 staining in CD, WD and refed groups of animals at weeks 48 (n=3 for animals). Data are shown as individual points (B, D, F, G and I). Statistics are based on one-way ANOVA (I) or two-way ANOVA (B, D, F and G), \**p*<0.05, \*\**p*<0.01, \*\*\**p*<0.001, \*\*\*\**p*<0.0001 and ns (\**p*>0.05).

## Discussion

Vascular cells are highly susceptible to increased oxidative stress elicited by disrupted energy metabolism, and the consequential damage in various vascular beds manifests as tissue-specific vascular diseases that are the leading cause of morbidity and mortality among individuals with metabolic disorders. While vessel abnormalities in insulin-sensitive tissues have been extensively registered over the years[29, 75], functional impacts of obesity on islet vessels remain relatively unappreciated and our knowledge of islet vessel pathology is largely limited to morphological characterizations. Remodeling of islet vasculature was consistently documented in several animal models of obesity[6, 7]. In the current study, we have described a time-resolved perturbation of islet vessel homeostasis in a WD-induced obese mouse model, marked by morphological remodeling, endothelial cell desensitization to VEGF-A due to impaired receptor internalization, compromised vascular barrier function as well as hemodynamic dysregulation in vivo. Moreover, we have also provided evidence for the deleterious effects of dysfunctional islet vessels on glucose homeostasis by hindering insulin outflow into the circulation.

In addition to the emergence of metabolic disorders in mice fed with WD, we detected significant morphological alterations and functional disorders in islet vessels by 12 weeks of WD feeding, which deteriorated with minimal recovery as the diet intervention continued. However, as islet vessels are not among the earliest responders to WD, there is a reasonable window for timely prophylaxis, which has significant implications for the prevention of chronic vascular complications related to obesity. Intriguingly, our investigations also revealed the heterogeneity in islet vascular cells under metabolic stress, evidenced by uneven islet vessel enlargement and varied Ca^2+^ response of individual capillary segments in WD-fed animals. Phenotypic and functional heterogeneity has been extensively documented among vascular beds in different organs[76, 77], and gene signatures in capillary endothelial cells exhibit a clear tissue-specific pattern[78, 79]. Further studies are required to decipher the molecular mechanisms behind vessel heterogeneity within the same islet at higher resolution to identify potential subpopulations of vascular cells that are more susceptible to this type of diet intervention.

Sustained islet vessel dysfunction was observed not only in WD-fed mice, but also in the refed mice. While removal of WD after 24 weeks and replacement with CD in these mice led to almost immediate alleviation of metabolic disorders, islet vasculature still resembled that of the WD group with enlarged diameters. More importantly, islet vessels in refed mice exhibited only partially regained fenestrae and slightly recovered permeability, while diminished intracellular Ca^2+^ mobilization in response to VEGF-A, increased basement membrane thickness as well as hemodynamic dysregulation persisted after 24 weeks of CD refeeding. Overall, none of the measured parameters were restored to the levels of the CD group by the end point, which points to poor vascular recovery and suggests a memory effect of the dietary stressors on islet vessels. The phenomenon that vascular endothelial cells retain cellular imprints generated by previous metabolic stress without the current presence of the stressor, also termed as metabolic memory, has been characterized in macro- and microvessels of type 2 diabetic patients[80–82]. A few pathways have been proposed as the molecular basis underlying the metabolic memory in endothelial cells. In our animal model, excessive dietary intake of saturated fatty acids and sucrose by continuous WD feeding may lead to the build-up of intracellular reactive oxygen species (ROS) and advanced glycation end products (AGEs), which can be generated from various intracellular sources and lead to increased oxidative stress, mitochondria damage as well as modifications of epigenetic signatures[83–85]. Considering the accumulative nature of this memory effect, our results highlight the necessity of prophylaxis and early intervention in preserving islet vessel function under pathological conditions such as obesity. It is thus likely that the reversibility of vessel pathogenesis would have been higher if we had conducted the diet switch at an earlier point during the time course. Additionally, in light of the known differences between genders in their response to diet manipulations[86, 87], it would be highly intriguing to investigate if female mice fed with WD display a higher level of islet vascular adaptation to the increased metabolic stress, and whether WD-induced VEGF-A desensitization and islet vessel dysfunction would be easier to reverse.

Anatomically interposed between the endocrine cells and the blood stream, islet vessels are not inert conduits of nutrients and oxygen. They actively coordinate nutrient sensing as well as hormone output of islets, and exhibit large plasticity under physiological conditions by quickly responding to biochemical stimuli in the microenvironment and adapting their barrier properties as well as vascular tone to islet metabolic activity[88, 89]. This coupling may be disrupted in case of obesity and metabolic disorders, due to the morphological modifications and functional impairments in islet vessels. Indeed, while the β cell secretory capacity in 12-week WD fed mice was proven to be intact in an ex vivo perifusion experiment, dysfunctional islet vessels impeded the outflow of secreted insulin into the blood stream when the animals received an intravenous glucose bolus. In comparison to intraperitoneal glucose tolerance tests, direct intravenous administration of glucose ensures better controlled delivery of glucose to islets without the interference of other internal organs, thereby allowing us to reveal the importance of islet vascular function in glucose handling. The amount of secreted insulin that crossed the vascular barrier to reach the blood stream during the first 3 min was significantly less in WD fed mice as compared to the CD fed mice, in spite of slightly higher islet vascular density in the former group. Therefore, islet vessel dysfunction is a prominent element in the manifestation of glucose intolerance in WD fed animals. Moreover, when the second cohort of mice was subjected to similar challenges of glucose bolus, the lean refed mice still displayed mild glucose intolerance despite normalized body weight and insulin sensitivity. The plasma insulin peak in refed mice remained lower than that of the control mice, although their insulin secretion capability was indistinguishable when evaluated in the perifusion system. Therefore, persisted islet vessel dysfunction in refed mice continued to hinder islet insulin outflow, and it played a causal role in the modest but significant delay in glucose clearance when the other risk factors were eliminated. Together, these results indicate that intact vascular barrier is crucial for insulin outflow, and disrupted islet vessel function directly undermines glucose homeostasis. If not restored during the early phase of pathogenesis, it may exacerbate the metabolic outcomes at later stages such as β cell failure in diabetes.

VEGF-A/VEGFR2 signaling is a key modulator of islet vascular morphology and function. Both hypo- and hypervascularization of islets caused by manipulation of islet VEGF-A expression level lead to disturbances in glucose metabolism[14, 16, 17]. It is noteworthy that not only islet endocrine cells are the main sources of pancreatic VEGF-A production, we also detected a higher expression level of VEGFR2 in islet vessels than the vessels of exocrine pancreas, which is consistent with previous documentation[14], and implies that islet vessel functionality may be highly susceptible to perturbations in VEGF-A/VEGFR2 signaling. Hence, it is crucial for islet homeostasis that both the expression of the VEGF-A ligand and the activity of this paracrine signal remain at a physiological level. In the current study, we showed that the level of gene expression, and more importantly, intra-islet VEGF-A/VEGFR2 signaling activity in vivo was modified by WD feeding, which was associated with pathogenesis of islet vessel dysfunction. Islet expression and production of VEGF-A was increased in WD-fed mice after 8 to 12 weeks of diet intervention, whereas endothelial cell responsiveness to VEGF-A declined almost simultaneously. Moreover, the lack of VEGF-A sensitivity persisted after 24 weeks of refeeding, although islet expression and production of VEGF-A was lowered to the control levels. Meanwhile, plasma level of the natural VEGF-A trap sFlt-1 was not altered by diet intervention during the entire course, indicating that ligand accessibility is unlikely the limiting factor for the strength of intra-islet VEGF-A signal activity. On the other hand, gene expression analysis and immunofluorescence staining confirmed the abundance of VEGFR2 on islet endothelial cell membrane during WD intervention, suggesting that not the expression level of the receptor, but its signaling activity may be compromised by continuous WD feeding, which leads to inadequate downstream signal transduction in endothelial cells. Recent studies have established the role of aPKCs in the regulation of VEGFR2 signaling by inhibiting its internalization[74]. The PKC family kinases are known to be involved in cellular stress responses, due to their general sensitivity to redox perturbations[90]. aPKCs, namely PKCζ and PKCι/λ, are unique members of this family, since their activation does not depend on diacylglycerol or Ca^2+^ like the other isoforms. Instead, their localization and activities in various cell types are regulated by anionic phospholipids and sphingolipids, such as phosphatidylinositol, phosphatidic acid and ceramide[91–94]. Prolonged high dietary fat intake by western diet feeding may lead to accumulation of these lipids and alteration of the membrane lipid profile[95, 96], which in turn give rise to over-activation of aPKC in islet endothelial cells. Indeed, we didn’t observe significant reduction in membrane-bound VEGFR2 by incubation with Pal/D-glu simultaneously, which was described in glucose treated cells in a previously mentioned study[36]. This discrepancy is probably due to different experimental conditions and endothelial cell types. Instead, we noted a markedly higher baseline level of aPKC phosphorylation and decreased VEGFR2 internalization upon activation in HUVECs treated with Pal/D-glu compared to control cells, which is accompanied by significantly reduced activation AKT and ERK1/2. Interestingly, high baseline aPKC phosphorylation was also observed in the recovered HUVECs, and VEGFR2 downstream signaling remained blunted as well, providing a possible explanation to the irreversibility of endothelial desensitization to VEGF-A in vivo. As aPKC activation by lipids is usually acute[91, 97], it is likely that after previous exposure to Pal/D-glu, lipid abnormities and potential modifications of other upstream signals persisted and continued to activate aPKC in the recovered HUVECs. Nevertheless, as the whole picture of aPKC activity regulation in endothelial cells remain incompletely understood, further investigations are required to unravel the exact signaling components that are sensitive to WD treatment for the development of potential therapeutic strategies aimed at vascular protection.

Besides the high dietary intake of sugar and fat, other risk factors may also account for islet vessel dysfunction in our model of long-term WD intervention. For example, ageing is related to endothelial cell senescence and dysfunction, and it also has a profound effect on VEGF-A signaling activity. A recent study found reduced VEGF-A signaling activity in multiple key organs of aged mice due to an increased production of sFlt-1[72]. In our study, we also observed a small and time-dependent decline in VEGF-A–triggered Ca^2+^ mobilization in the CD-fed mice, which may be due to ageing. Therefore, ageing probably acted synergistically with metabolic stress in the development of islet vascular endothelial cell VEGF-A desensitization and vessel dysfunction in WD-fed mice. Future studies using our versatile transplantation platform will allow isolating the ageing factor and further investigating the sole impacts of vascular ageing on islet function, by mismatching the ages of islet donors and recipients, e.g., by transplanting old animals with islets derived from young donors, which will become re-vascularized by aged endothelial cells.

While we focused on alterations in islet endothelial VEGF-A signaling activity and related vessel function, it is worthwhile to evaluate the plasticity and heterogeneity of other vascular cell types under WD feeding to gain a complete overview of its vascular impacts. Islet pericytes, for instance, may play a role in the islet vessel stiffness and ultrastructural modifications observed in WD-fed mice, as they constitute another element in the fine regulation of local blood flow, and also contribute to the deposition of certain matrix proteins in the islet basement membrane[57, 98].

In summary, we have delineated the time-resolved impairment of islet vascular function as well as its metabolic consequences in a WD-induced obesity animal model, and identified intra-islet VEGF-A/VEGFR2 signaling obstruction as an underlying mechanism. By scrutinizing the reversibility of islet vascular defects caused by WD feeding, we highlight the role of islet vascular dysfunction in the manifestation of diabetic phenotypes under pathological conditions such as obesity, and emphasize the necessity in retaining physiological activity of intra-islet VEGF-A signaling for optimal islet function and maintenance of glucose homeostasis.

## Supporting information

Xiong et al Supplemental materials

## Acknowledgements

We thank Lars Haag at Karolinska Institutet’s core facility for electron microscopy for obtaining the TEM images, and Dr. Stefan Jacob for help with image analysis.

## Sources of funding

This work was supported by funding from Karolinska Institutet, the Strategic Research Program in Diabetes at Karolinska Institutet, the Swedish Research Council, the Novo Nordisk Foundation, the Swedish Diabetes Association, the Family Knut and Alice Wallenberg Foundation, the Jonas & Christina af Jochnick Foundation, the Family Erling-Persson Foundation, Berth von Kantzow’s Foundation and European Research Council grant ERC-2018-AdG 834860 EYELETS.

## Author contributions

Y.X. coordinated and designed the study. Y.X, E.I. and P.-O.B. conceived the study. E.I. and P.-O.B. supervised the study. Y.X., A.D., M.V. and E.I. performed and analyzed the experiments. Y.X. wrote the manuscript. E.I. and P.-O.B. revised and edited the manuscript. All authors reviewed the results and approved the final version of the manuscript.

## Disclosures

P.-O.B. is the founder and CEO of Biocrine AB.

## Supplemental Material

Tables S1

Figure S1-S5

Videos S1–S3

## Notes

### Summary of Updates

Supplemental materials are added.

