## Supplementary material for "Diet-induced obesity results in endothelial cell desensitization to VEGF-A and permanent islet vascular dysfunction": Xiong et al Supplemental materials

**Supplemental Table S1**. Sequences of qPCR primers used in this study.

| Gene | Sequence |
| --- | --- |
| *Tbp* | L:5’-TGC TGT TGG TGA TTG TTG GT -3’<br>R:5’-CTG GCT TGT GTG GGA AAG AT -3’ |
| *Vegfa* | L:5’-CAG GCT GCT GTA ACG ATG AA -3’<br>R:5’-GCA TTC ACA TCT GCT GTG CT -3’ |
| *Kdr* | L:5’-GGC GGT GGT GAC AGT ATC TT -3’<br>R:5’-GTC ACT GAC AGA GGC GAT GA -3’ |
| *Pecam1* | L:5’-CTG AGC CTA GTG TGG AAG GC -3’<br>R:5’-GTC TCT GTG GCT CTC GTT CC -3’ |
| *L19* | L:5’-GGC ACA TGG GCA TAG GTA AG -3’<br>R:5’-CCA TGA GAA TCC GCT TGT TT -3’ |
| *KDR* | L:5’-AAC GGC GCT TGG ACA GCA -3’<br>R:5’-CAT GCC CTT AGC CAC TTG GAA -3’ |
| *EFNB2* | L:5’-GTA CCG GAG GAG ACA CAG GA -3’<br>R:5’-CCT TCT CGT AGT GAG GGC AG -3’ |
| *PARD3* | L:5’-CAG CCT TCC CAC TCT CTG GA -3’<br>R:5’-TTC TCG GGC TTC AGT TTG GC -3’ |
| *DAB2* | L:5’-CGG GTG TTG ACC AGA TGG ATT -3’<br>R:5’-AAC AGG AGG TGA TGC CGT TT -3’ |

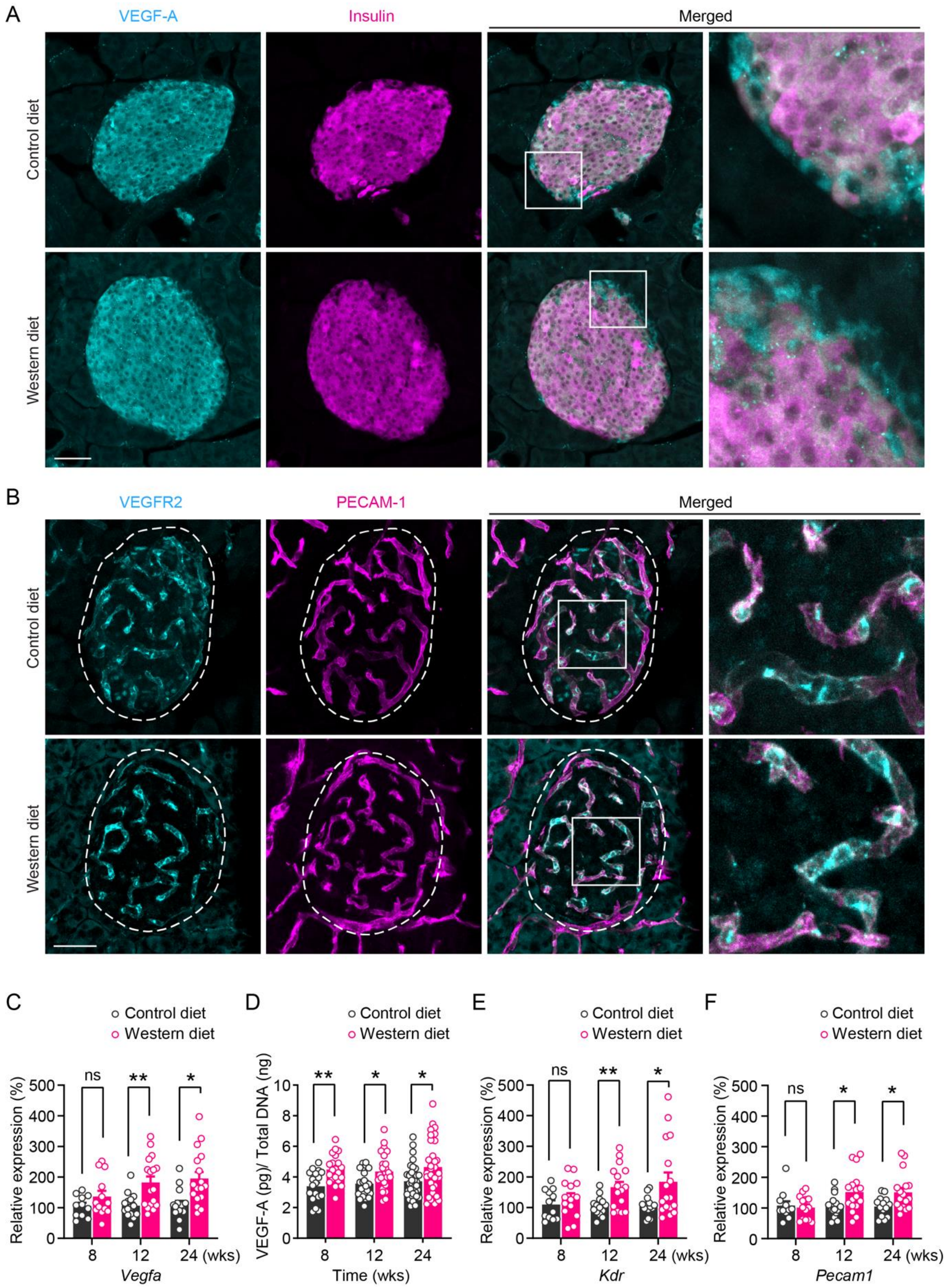


**Figure S1**. Western diet enhances islet expression and production of VEGF-A.

**A-B**, Representative confocal images are presented as maximum intensity projections, showing immunofluorescence staining of pancreatic sections from CD- and WD-fed animals at week 8 of diet intervention, showing VEGF-A (cyan), insulin (magenta) (A), VEGFR2 (cyan) and PECAM-1 (magenta) (B). Squares indicate areas magnified in the right panels. Scale bars: 50 μm. **C,** Gene expression level of *Vegfa* in freshly isolated islets from CD- and WD-fed animals at weeks 8, 12 and 24 of diet intervention (CD: n=11, 15, 15; WD: n=11, 16, 15). **D,** Ex vivo VEGF-A production in islets from CD- and WD-fed animals at weeks 8, 12 and 24 of diet intervention (CD: n=6, 6, 8; WD: n=6, 8, 8). **E-F**, Relative gene expression levels of *Kdr* (E) and *Pecam1* (F) in freshly isolated islets from CD- and WD-fed animals at weeks 8, 12 and 24 of diet intervention (CD: *Kdr*, n=12, 14, 15; *Pecam1*, n=11, 15, 14; WD: *Kdr*, n=14, 16, 17; *Pecam1*, n=14, 17, 17). Data are shown as individual points (C-F). Statistics are based on multiple unpaired *t*-tests (C-F), *$p<0.05$, **$p<0.01$ and ns (*$p>0.05$).

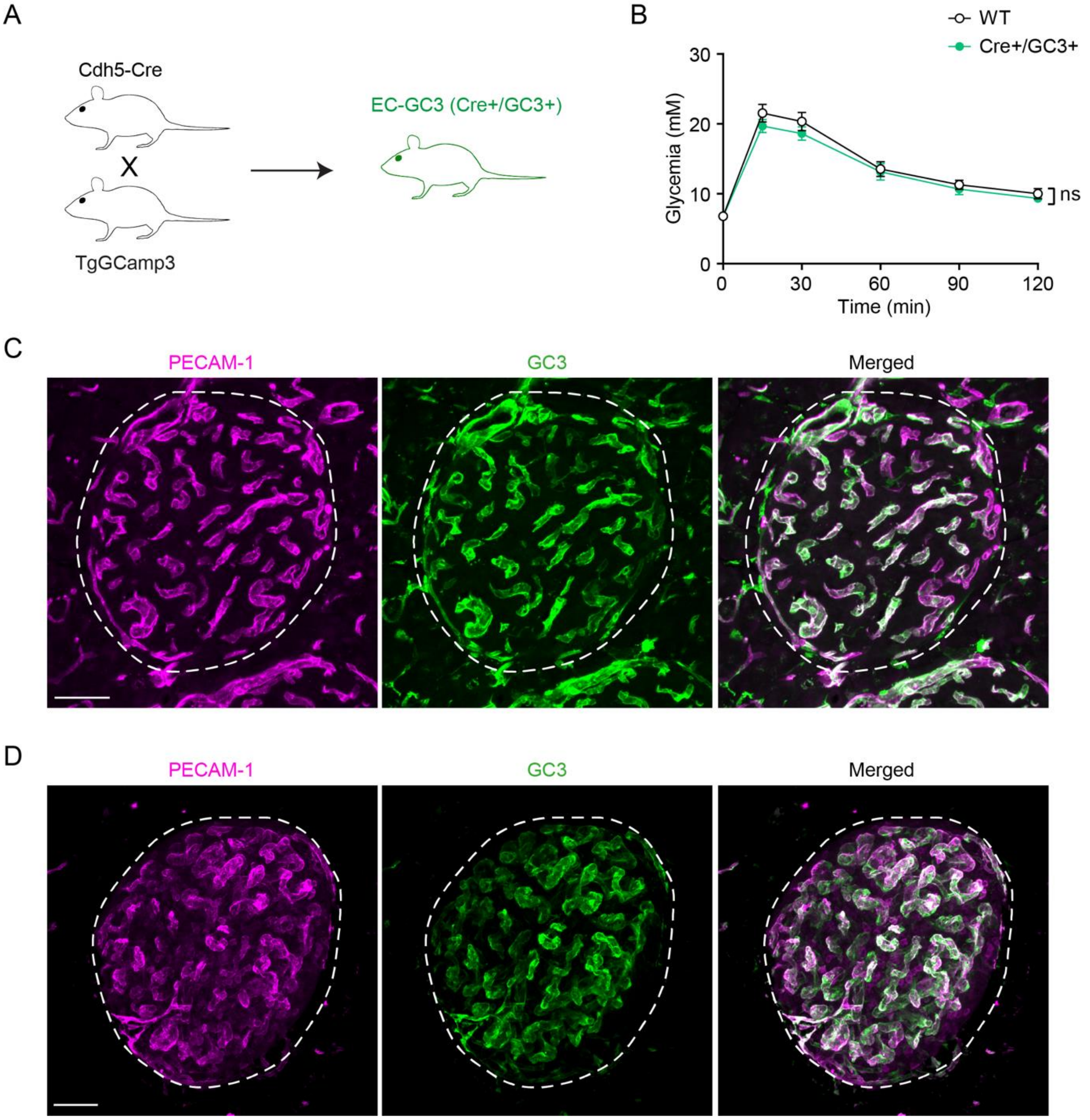


**Figure S2**. Characterization of the EC-GC3 reporter mouse line.

**A**, Generation of the EC-GC3 reporter mouse line. **B**, i.p.GTT of EC-GC3 (n=14) and their wild type littermates (WT, n=15). Data are shown as mean ± SEM. Statistics are based on two-way ANOVA, ns (*$p$>0.05). **C-D,** Representative confocal images are presented as maximum intensity projections, showing immunofluorescence staining of pancreatic sections (C) and whole-mount islet grafts (D) dissected from EC-GC3 animals, showing PECAM-1 (magenta) and GC3 (GFP, green). Scale bars: 50 μm.

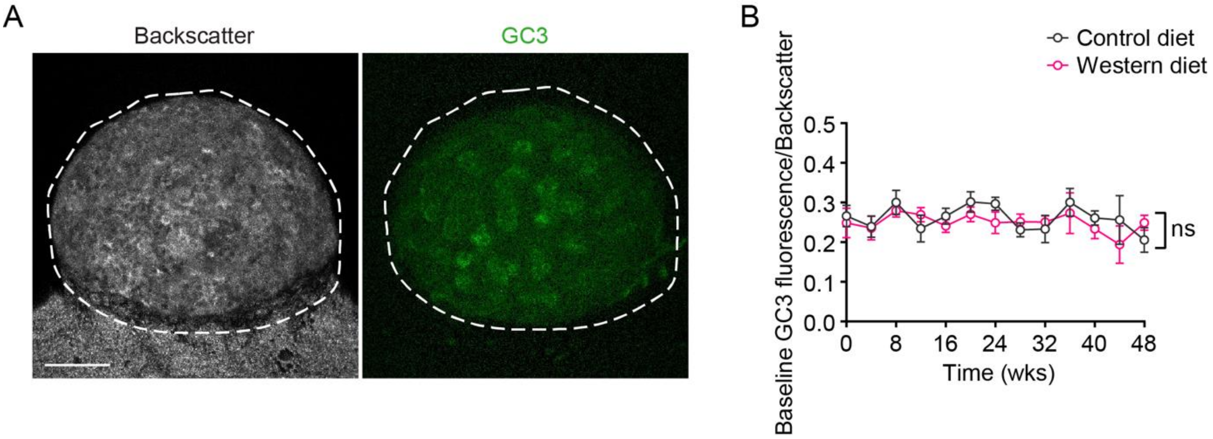


**Figure S3**. Baseline GC3 fluorescence remains stable during diet intervention.

**A,** Representative confocal images are presented as maximum intensity projections, showing backscatter signal (left) and GC3 fluorescence (right) in an islet graft prior to intravenous VEGF-A injection. Scale bar: 50 µm. **B,** Normalized baseline GC3 fluorescence fluctuations over diet intervention in CD- (n=8-11) and WD-fed (n=8-15) animals. Data are shown as mean ± SEM. Statistics are based on two-way ANOVA, ns (*$p$>0.05).

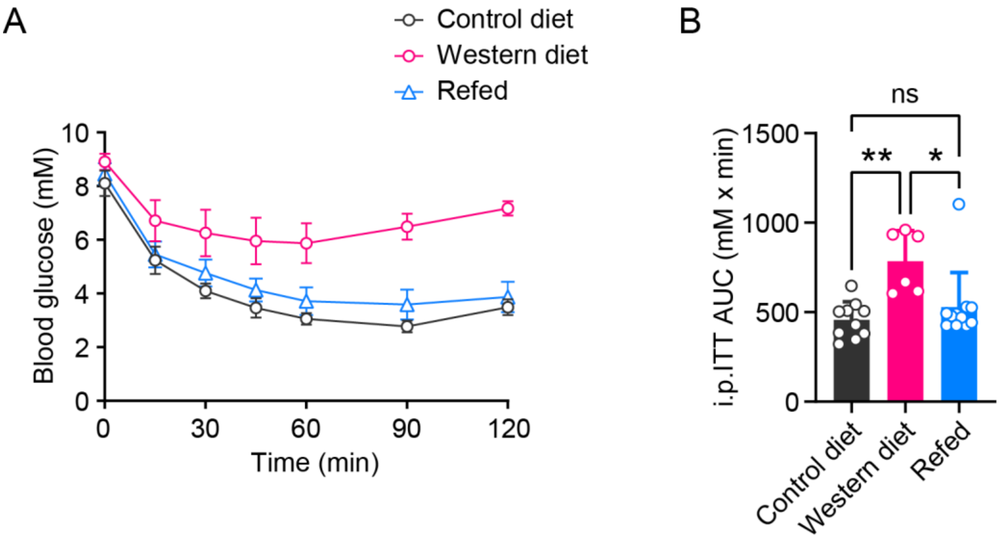


**Figure S4**. Insulin sensitivity recovers in refed animals.

**A**, Intraperitoneal insulin tolerance test (i.p.ITT) in CD (n=10), WD (n=6) and refed (n=11) groups of animals at week 48. **B,** Area under the curve for insulin tolerance test. Data are shown as mean ± SEM (A) or individual points (B). Statistics are based on one-way ANOVA (B), **p*<0.05, ***p*<0.01, ns (**p*>0.05).

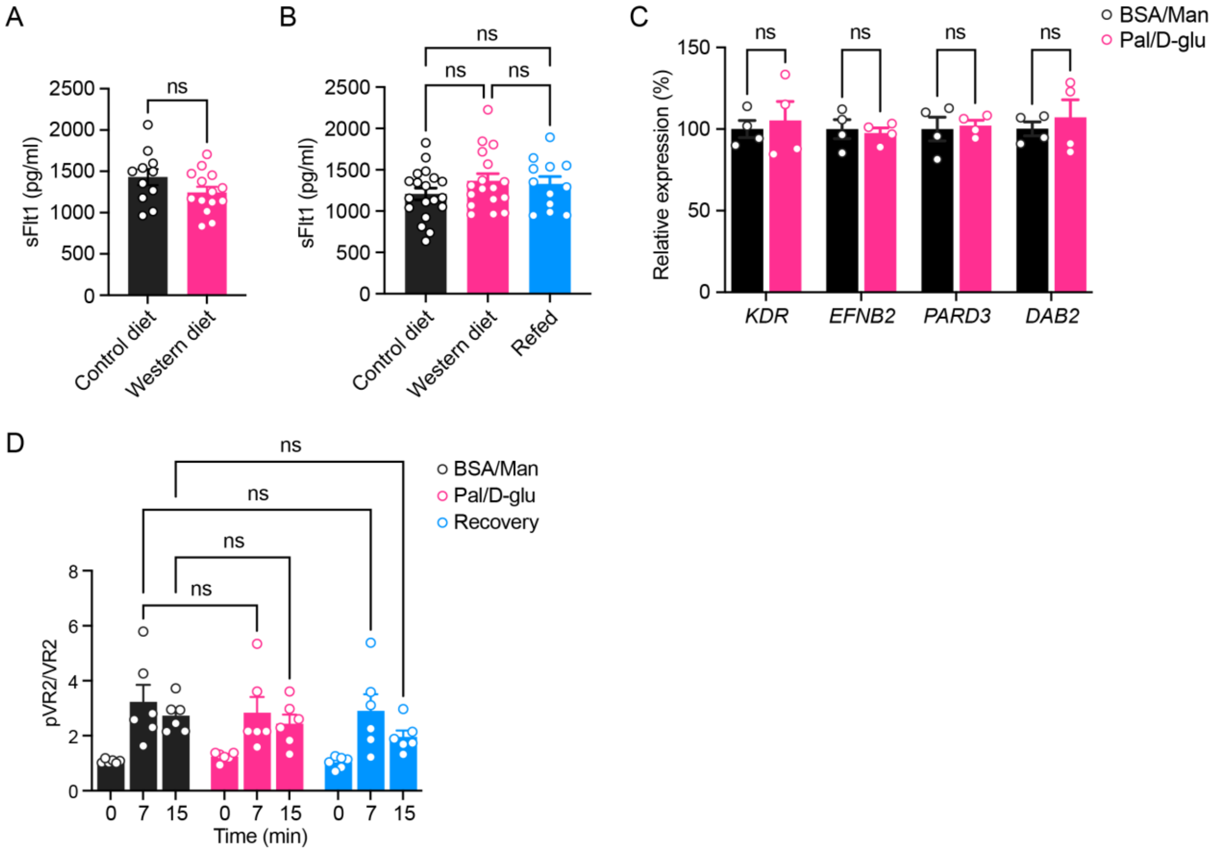


**Figure S5.** Several factors involved in the regulation of VEGFR2 signaling activity remain unchanged by western diet elements.

**A,** Plasma sFlt-1 levels in CD (n=11) and WD (n=14) groups of animals at week 24. **B,** Plasma sFlt-1 levels in CD (n=19), WD (n=17) and refed (n=12) groups of animals at week 48. **C,** Relative gene expression levels of *KDR* and components of VEGFR2 sorting complex in HUVECs cultivated under BSA/Man or Pal/D-glu conditions (n=4). **D,** Normalized VEGFR2 Tyr1175 phosphorylation levels during VEGF-A stimulation in HUVECs cultivated under BSA/Man, Pal/D-glu or recovery conditions (n=4). Data are shown as individual points (A to D). Statistics are based on unpaired t-tests (A and C), one-way ANOVA (B) and two-way ANOVA (d). ns (*$p$>0.05).

**Supplemental Video S1**. Representative movie showing intra-islet vessel $Ca^{2+}$ response to VEGF-A bolus in CD-fed animals at week 12. Time series of confocal images are presented as maximum intensity projections. The movie is accelerated, and real time duration is 8 min. VEGF-A was injected at 1 min as indicated. Scale bar: 50 μm.

**Supplemental Video S2**. Representative movie showing intra-islet vessel $Ca^{2+}$ response to VEGF-A bolus in WD-fed animals at week 12. Time series of confocal images are presented as maximum intensity projections. The movie is accelerated, and real time duration is 8 min. VEGF-A was injected at 1 min as indicated. Scale bar: 50 μm.

**Supplemental Video S3**. Representative movie showing the labeling of RBCs for in vivo quantification of islet blood flow velocity in a control animal. Scale bar: 50 μm.
